# The PPR domain of mitochondrial RNA polymerase is a ribonuclease required for mtDNA replication

**DOI:** 10.1101/2021.03.12.435139

**Authors:** Yi Liu, Zhe Chen, Zong-Heng Wang, Katherine Delaney, Juanjie Tang, Mehdi Pirooznia, Duck-Yeon Lee, Yuesheng Li, Hong Xu

## Abstract

Mitochondrial DNA (mtDNA) replication and transcription are of paramount importance to cellular energy metabolism. Mitochondrial RNA polymerase (POLRMT) is thought to be the primase for mtDNA replication. However, it is unclear how POLRMT, which normally transcribes long polycistronic RNAs, can produce short RNA oligos to initiate mtDNA replication. Here we show that the PPR domain of *Drosophila* POLRMT is a 3’ to 5’ exoribonuclease. The exoribonuclease activity is indispensable for POLRMT to synthesize short RNA oligos and to prime DNA replication *in vitro*. An exoribonuclease deficient POLRMT, POLRMT^E423P^ partially restores mitochondrial transcription but fails to support mtDNA replication when expressed in *POLRMT* mutant background, indicating that the exoribonuclease activity is necessary for mtDNA replication. Overexpression of POLRMT^E423P^ in adult flies leads to severe neuromuscular defects and a marked increase of mtDNA transcripts errors, suggesting that exoribonuclease activity may contribute to the proofreading of mtDNA transcription. PPR domain of human POLRMT also has exoribonuclease activity, indicating evolutionarily conserved roles of PPR domain in mitochondrial DNA and RNA metabolism.

## Introduction

Mitochondria, the energy-harnessing organelles of eukaryotic cells, carry out oxidative phosphorylation that produces more than 90% of cellular ATP under aerobic conditions (Noll et al., 1990). Mitochondria contain their own genome, the mitochondrial DNA (mtDNA) inside the matrix (Wallace, 2005). Mammalian mtDNA is a double-stranded circular DNA molecule, encoding 13 core components of oxidative phosphorylation system, 2 rRNAs and 22 tRNAs required for the synthesis of mtDNA-encoded proteins (Wallace, 2005). Dysregulations of mtDNA maintenance and mitochondrial genes expression often impair cellular energy metabolism and have been associated with a broad spectrum of human diseases (Wallace, 2005).

Animal mtDNA is highly compact and usually lacks introns or regulatory sequences, aside from a large non-coding region containing promoters for both heavy (H) and light (L) strands and the replication origin of H-strand (Bonawitz et al., 2006). According to the strand displacement model, mtDNA replication is initiated at the H-strand origin and proceeds unidirectionally for about two-thirds of the circular genome, which exposes the origin of L-strand and allows the replication of L-strand in the opposite direction (Gustafsson et al., 2016). Besides this asymmetric replication model, other mechanisms, in which both strands are synthesized synchronously are also demonstrated (Holt et al., 2000; Joers and Jacobs, 2013). MtDNA replication is carried out by DNA polymerase γ (pol γ) (Falkenberg et al., 2007). However, pol γ cannot initiate mtDNA replication from scratch. Instead, short RNA oligos are required to prime the DNA replication (Kornberg, 1984), particularly in models of synchronous replication. Currently, the molecular identity of mitochondrial primase remains unsettled. A primase-polymerase (PrimPol) that involves in nuclear genome replication and repair has been reported to be co-sedimented with mitochondria and proposed to be a primase for mtDNA replication (García-Gómez et al., 2013). Yet, PrimPol is found in chordates only (García-Gómez et al., 2013), its prevalence and potential roles in mtDNA replication in other animal phyla remain to be explored. The mitochondrial RNA polymerase (POLRMT), a single subunit RNA polymerase carrying out mtDNA transcription is also required for mtDNA replication (Shadel and Clayton, 1997). The conditional knockout of POLRMT not only abolishes mtDNA transcription, but also severely reduces both steady-state mtDNA level and *de novo* mtDNA synthesis in mouse heart (Kühl et al., 2016). Additionally, POLRMT produces short RNA oligos that can be used to prime DNA synthesis *in vitro* (Wanrooij et al., 2008), suggesting that POLRMT could be the mitochondrial primase. However, it is not clear how POLRMT, which transcribes both strands of mtDNA to long polycistronic transcripts (Hillen et al., 2018), can switch from continuous transcription to primer synthesis.

Several protein coding genes on mtDNA are overlapping, and in a few cases, part of the stop codons are missing (Wolstenholme, 1992). Additionally, all mitochondrial mRNAs and rRNAs are polyadenylated (Mercer et al., 2011). Thus, the primary polycistronic transcripts have to be properly processed to produce mature RNA species (Ojala et al., 1981). Consistent with this notion, many RNA modifying enzymes and RNA binding proteins are found inside mitochondria (Holzmann et al., 2008; Sanchez et al., 2011). The pentatricopeptide repeat (PPR) proteins have emerged as major modulators of post-transcriptional regulation in organelles (Lurin et al., 2004). PPR is a degenerate 35 amino acid motif repeated in tandem, ranging from 2 to over 20 (Schmitzlinneweber and Small, 2008). To date, almost all annotated PPR proteins are predicted to localize in either chloroplasts or mitochondria (Rovira and Smith, 2019). PPR proteins are characterized as RNA-binding proteins and suggested to regulate organellar transcription, RNA processing, stability control and translation regulations (Manna, 2015). PPR proteins are found in all eukaryotes, but the number of PPR proteins has remarkably expanded in land plants (Small and Peeters, 2000), which generally have large organellar genomes (Fujii and Small, 2011), further substantiating their roles in organellar RNA metabolism.

*Drosophila melanogaster* is an excellent model to study mtDNA genetics and gene expression. Between flies and mammals, mitochondrial gene contents are identical, structure and gene organization of mtDNA are similar, and most nuclear factors involved in mtDNA maintenance and mitochondrial RNA metabolism are highly conserved (Sánchez-Martínez et al., 2006; Wolstenholme, 1992). Moreover, the short life cycle and relatively simple anatomy make *Drosophila* particularly amendable to explore the developmental and physiological significance of mitochondrial regulations. There are only 5 and 7 PPR genes in fly and human nuclear genomes, respectively (Baggio et al., 2014; Lightowlers and Chrzanowska-Lightowlers, 2013). All of these proteins are either experimentally proved or annotated to be localized in mitochondria (Lightowlers and Chrzanowska-Lightowlers, 2013; Manna, 2015). Leucine-rich PPR-containing protein (LRPPRC) regulates mitochondrial RNA processing (Harmel et al., 2013). Mitochondrial ribosomal protein of the small subunit 27 (MRPS27) is involved in regulation of mitochondrial translation (Davies et al., 2012). Mitochondrial RNase P protein 3 (MRPP3), a subunit of the major enzyme complex responsible for mitochondrial endonucleolytic cleavage, is required for tRNA processing (Holzmann et al., 2008). A cluster of PPR repeat domain proteins, PTCD1, PTCD2 and PTCD3 are involved in mitochondrial transcripts processing and translation (Rackham et al., 2009; Sanchez et al., 2011; Xu et al., 2008). Nonetheless, the biochemical basis of PPR proteins in RNA processing remains unclear. POLRMT is also a PPR protein. It has two PPR motifs at the N terminus, besides the C-terminal polymerase domain that is highly homologous to phage T7 RNA polymerase. Structural analyses reveal that the PPR domain of POLRMT locates at the exit channel of newly synthesized RNA (Ringel et al., 2011). However, the exact molecular function of PPR domain in transcriptional regulation is currently unknown.

Here we report that PPR domain in *Drosophila* POLRMT is a 3’ to 5’exoribonuclease and is indispensable for the synthesis of short RNA oligos *in vitro* and mtDNA replication in tissues. Overexpression of an exoribonuclease deficient POLRMT in adult flies leads to severe premature aging phenotypes including impaired mobility, increased susceptibility to mechanical stress and reduced life span. While the level of mitochondrial transcripts and their processing are normal in the overexpressing flies, the frequency of transcription errors is markedly increased compared to control flies, indicating that the 3’ to 5’ exoribonuclease activity of POLRMT may contribute to mtDNA transcription proofreading. We find the PPR domain of human POLRMT also possesses 3’ to 5’ exoribonuclease activity, suggesting conserved roles of PPR domain in mtDNA replication and transcription.

## Results

### PPR domain is required for POLRMT to synthesize short RNAs

There is no PrimPol or related family members in *Drosophila* genome (https://www.ncbi.nlm.nih.gov/homologene/?term=primpol). In addition, human PrimPol did not localize to the mitochondrial matrix (Figure S1), based on a GFP complementation assay (Zhang et al., 2015), disputing the idea that PrimPol is the mitochondrial primase. Previous studies suggest that mammalian POLRMT could be the mitochondrial primase (Kühl et al., 2016; Wanrooij et al., 2008). To explore potential contribution of *Drosophila* POLRMT to mtDNA replication, we deleted the *CG4644* locus that encodes POLRMT by CRISPR mediated recombination. Homozygous mutant flies developed into 3^rd^ instar larva but were much smaller than wild type (Figure 1A). They were stuck at the 3^rd^ instar larval stage, and eventually died after 10 days. We fed larva with 5-Ethynyl-2’-deoxyuridine (EdU) for 12 hours, and visualized EdU with Click chemistry to label DNA replication. In wild type fat body, many EdU puncta were co-localized with mitochondrial transcription factor A (TFAM) that marks mitochondrial nucleoids (Figure 1B). While TFAM puncta were present in *POLRMT^del^*, no EdU incorporation was detected (Figure 1B), illustrating that POLRMT is required for *de novo* mtDNA replication in *Drosophila*.

**Figure 1.**
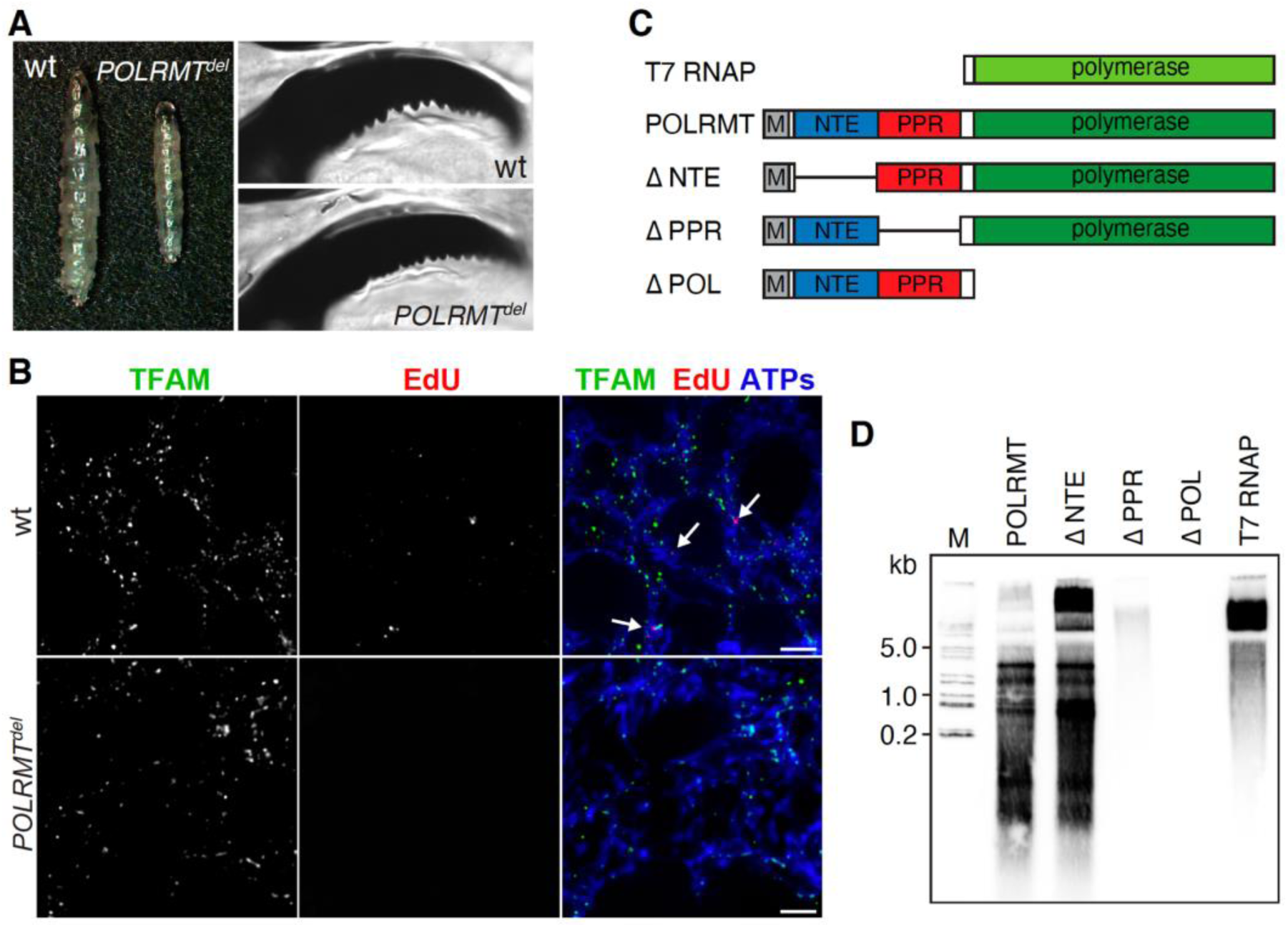
POLRMT is required for *de novo* mtDNA synthesis in *Drosophila*. (A) Comparison of body size and mouthpart between wild type (wt) and homozygous *POLRMT* knockout (*POLRMT^del^*) larvae. (B) Representative images of wild type (wt) and homozygous *POLRMT^del^* fat body labeled with 5-ethynyl-2’-deoxyuridine (EdU) and visualized with Alexa Fluor 555 dye to mark DNA replication. TFAM (Green) and ATP synthase (ATPs; Blue) were used to mark mtDNA and mitochondria, respectively. Arrows indicate the co-localization of TFAM and EdU signals. Scale bars, 10 µm. (C) Schematic diagram of T7 RNA polymerase, POLRMT protein and deletion mutants. Major domains are indicated. M, mitochondrial targeting signal (residues 1-76). NTE, N-terminal extension (residues 77-313). PPR, PPR domain (residues 314-464). Polymerase, the canonical polymerase domain (residues 511-1369). (D) RNA synthesis by a commercial T7 RNA polymerase (0.5 units) or recombinant proteins (500 fmol) on M13mp18 ssDNA template was analyzed on a denaturing polyacrylamide gel (10%). Molecular size markers are indicated.

To test whether *Drosophila* POLRMT can function as a primase, we examined its polymerase activity in an *in vitro* transcription assay using M13 phage genomic DNA as the template. Majority of POLRMT products were short RNA species less than 100 nt (Figure 1D). We included T7 RNA polymerase in parallel as a control, which mainly generated long RNA species (>1000 nt). Besides the polymerase domain that is highly homologous to T7 RNA polymerase, POLRMT also contains a largely flexible N-terminal extension (NTE) and two PPR motifs at its N-terminus (Figure 1C). To define which domain might alter the property of POLRMT from its evolutionary ancestor, T7 RNA polymerase, we expressed and purified a series of deletions of POLRMT and examined polymerase activity *in vitro* (Figure 1D). Deletion of NTE (ΔNTE) seemingly improved the overall polymerase activity (Figure 1D). Surprisingly, PPR domain deletion (ΔPPR) synthesized long RNA species (>1000 nt) only, which is similar to T7 RNA polymerase (Figure 1D). These results demonstrate that the PPR domain enables POLRMT to synthesize short RNA oligos.

### PPR domain in *Drosophila* POLRMT is a 3’-5’ exoribonuclease

Classical primases recognize short (usually 2-3 nt), degenerate single-stranded DNA (ssDNA) sequences to initiate the synthesis of RNA oligos (Griep, 1995). Given that PPR motifs have been proposed to be RNA binding proteins, we hypothesized that PPR might bind to some short DNA sequences to initiate RNA synthesis from these discrete sites on mtDNA. We expressed and purified the PPR domain of fly POLRMT from *E. coli* to near homogeneity (Figure S2A, B), and tested its affinity to a double-stranded DNA (dsDNA) fragment using electrophoresis mobility shift assay (EMSA). PPR domain did not appear to bind to dsDNA (Figure 2A). We then tested whether it might bind to DNA-RNA hybrid, as the PPR domain of POLRMT locates at the exit channel of newly synthesized RNA (Ringel et al., 2011). A potential PPR-DNA/RNA complex was detected in EMSA assay (Figure 2A). However, the overall intensity of probe was notably diminished in the presence of PPR (Figure 2A), which intrigued us to test whether the PPR domain may possess ribonuclease activity. Incubation of PPR with a 3’-Biotin-labeled single-stranded RNA (ssRNA) gradually decreased the intensity of full-length target RNA (Figure 2B), without the emergence of short, intermediate oligoribonucleotides that were evident in the reaction consisted of PPR with a 5’-Biotin-labelled ssRNA (Figure 2C). Additionally, the smallest cleavage products migrated about the same rate as Biotin-16-UTP, indicating that PPR has 3’ to 5’ exoribonuclease activity and generates mononucleotides (Figure 2D).

**Figure 2.**
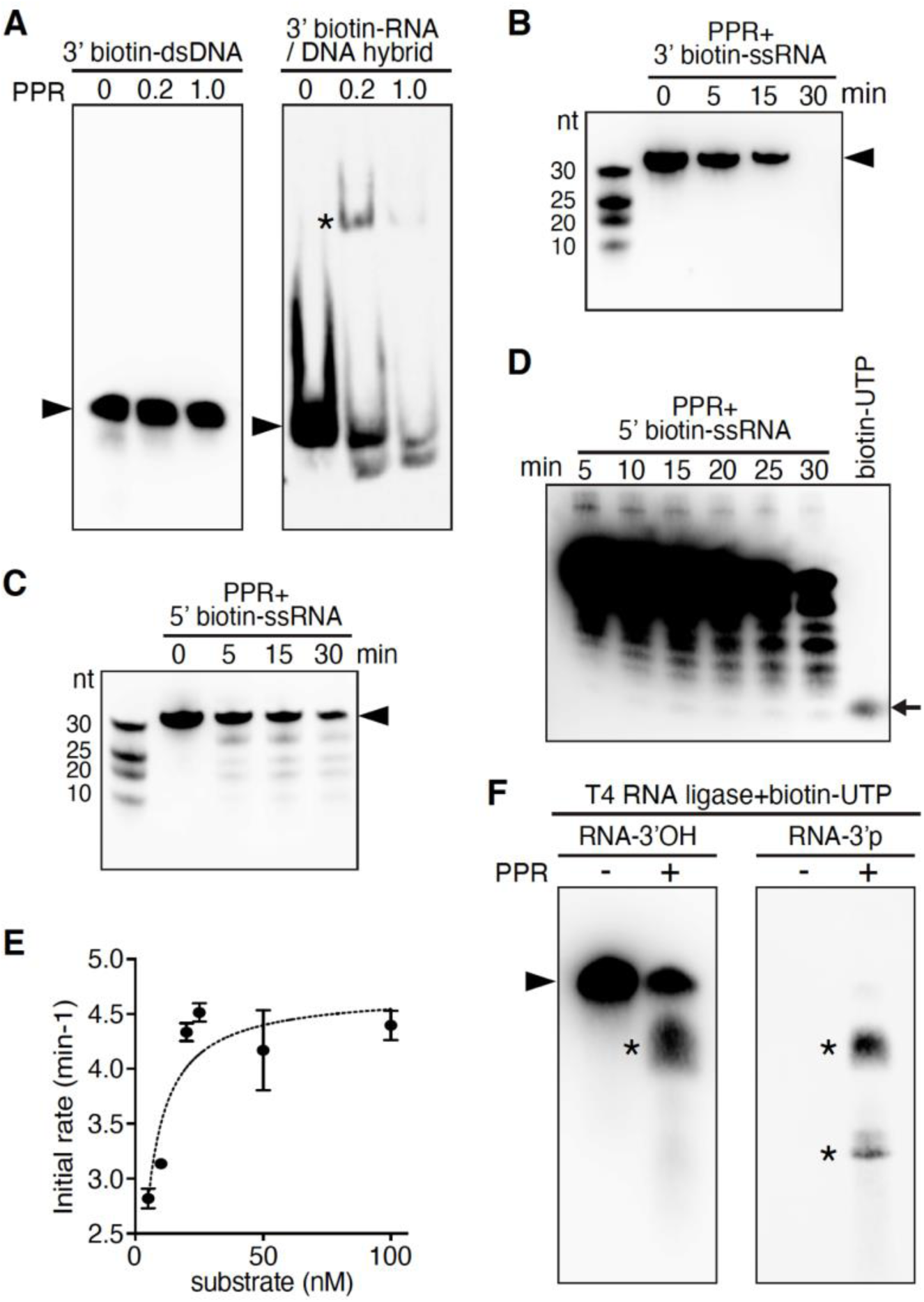
PPR domain in POLRMT is a 3’-5’ exoribonuclease. (A) Electrophoretic mobility shift assay of the recombinant PPR domain with 3’-Biotin-labeled dsDNA or RNA-DNA hybrid (50 nM). Concentrations (µM) of PPR domain in reactions are indicated. Asterisk: the RNA-DNA hybrid-protein complex; arrowhead: free probe. (B) PPR protein degrades ssRNA. 3’-Biotin-labeled target ssRNA (0.05 µM) was incubated with PPR protein in 10 µl reaction mix for indicated time (min) at 32 °C and separated on a denaturing polyacrylamide gel (20%). Arrowhead indicates the full-length RNA. (C) A same experiment as in (B) using 5’-Biotin-labeled ssRNA as the substrate. (D) A long-exposure image of same reaction mixes as in (C) after 5, 10, 15, 20, 25 and 30 min of incubation. Biotin-16-UTP (0.1 mM) was included to indicate the size of a biotinylated mononucleotide. The arrow indicates the end products of PPR protein degradation. (E) The Michaelis-Menten graph showing the kinetics of RNA cleavage by PPR protein using ssRNA as the substrate. *K_m_* = 3.421 nM, *K_cat_* = V_max_/E_0_ = 8.54 min^−1^. MW= 18166.9 g/mol. Each data point represents the average of three independent experiments and error bars represent sd. (F) The 3’ end of RNA cleavage products by PPR protein is hydroxyl group. 3’-hydroxyl ssRNA or 3’-phosphate ssRNA (0.5 µM) was preincubated with or without PPR protein (5 µM) in 10 µl reaction mix for 10 min at 32 °C, and the RNA products were labeled a single biotinylated (bis) phosphate to the 3’ terminus by T4 RNA ligase. Asterisk: intermediate RNA products; arrowhead: the full-length target RNA. Note that the full-length probe with 3’-hydroxyl group, not the one with 3’-phosphate group can be labelled with the biotinylated cytidine (bis) phosphate nucleotide, whereas degradation products of both probes can also be labelled.

We concerned that the detected ribonuclease activity might be resulted from RNase contamination in *E. coli* extracts. We have used polyethyleneimine precipitation to remove nucleic acids and nucleic acids binding proteins from the lysates (Cordes et al., 1990), to minimize the potential contamination. We also found that the ribonuclease activity of PPR required divalent metal ions, especially Mn^2+^ and Mg^2+^, and was sensitive to cation chelator (Figure S2C). On the contrary, common RNases in contaminations are resistant to metal chelating agents (Fersht, 1977). We determined the *k_m_* and *k_cat_* of PPR in a ssRNA degradation reaction to be 3.421 nM and 8.54 min^−1^, respectively (Figure 2E), demonstrating that PPR has relatively high-affinity to RNA substrate. However, its catalytic efficiency (*k_cat_/k_m_* = 4.16 × 10^7^ M^−1^s^−1^) is much lower than that of RNase A (Park and Raines, 2003), the most common RNase contamination. Furthermore, purified PPR protein was insensitive to broad spectrum RNase inhibitors, which at the same concentration, completely inhibited RNase A (Figure S2D). Last and most importantly, while most common RNases contamination generate 3’ phosphate group, the 3’ end of PPR degradation product was hydroxyl group, which can be conjugated with biotinylated cytidine by T4 RNA ligase (Figure 2F). Noteworthy, the full-length fly POLRMT protein also showed 3’-5’ exoribonuclease in the *in vitro* assay (Figure 3A, 3B). Taken together, these observations confirm that PPR domain of POLRMT is in fact a 3’-5’ exoribonuclease.

**Figure 3.**
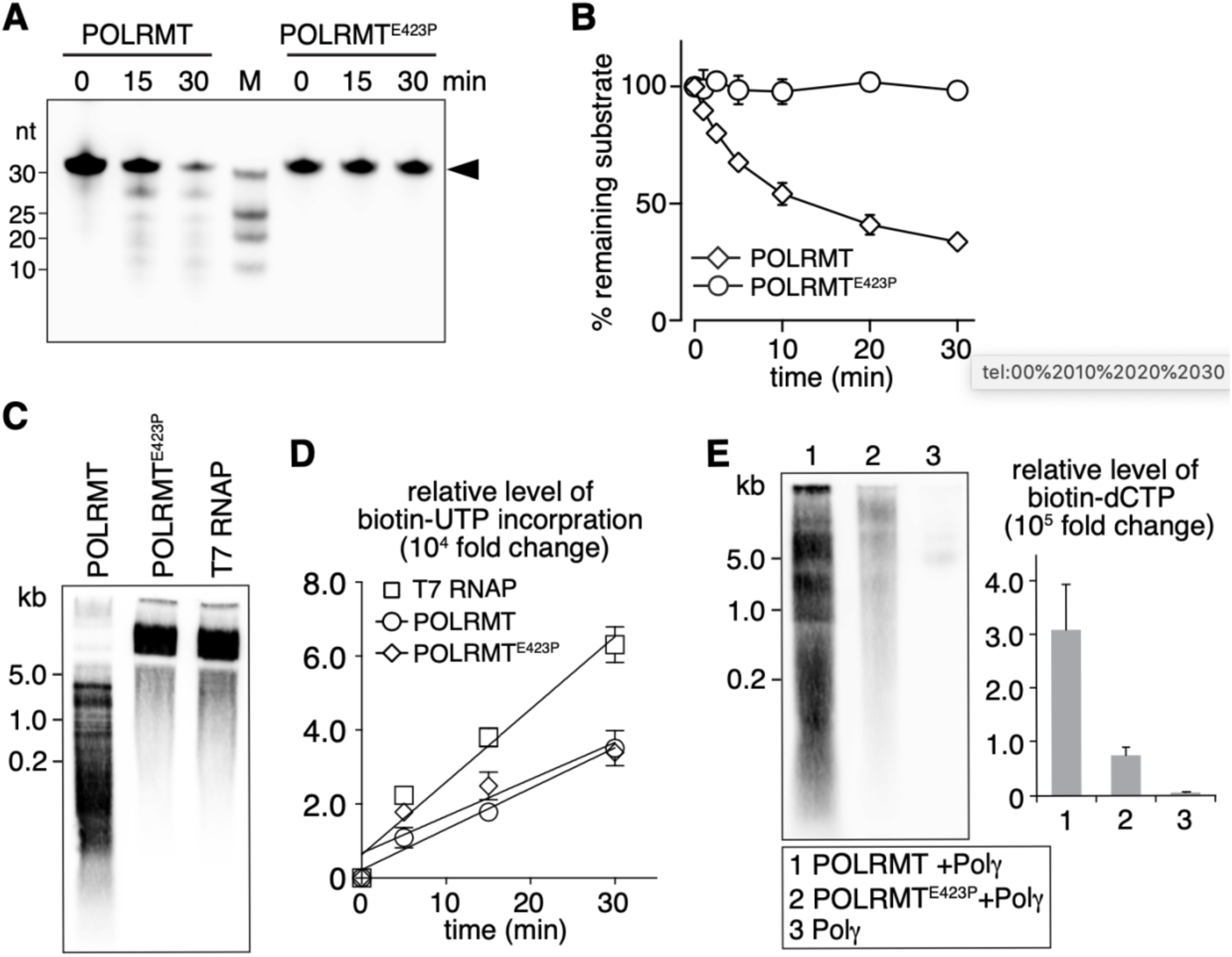
Exoribonuclease activity of POLRMT is essential for priming DNA replication *in vitro*. (A) POLRMT^E423P^ lacks the exoribonuclease activity. 5’-Biotin-labeled target ssRNA (0.05 µM) was incubated with recombinant POLRMT or POLRMT^E423P^ (0.5 µM) in 10 µl reaction mix for indicated time (min) and separated on a denaturing polyacrylamide gel (20%). Arrowhead indicates the full-length target RNA. (B) Pseudo-first-order cleavage kinetics of 5’-Biotin-labeled target ssRNA by POLRMT and POLRMT^E423P^. The level of remaining RNA in each time point was measured, normalized to the initial level and plotted. Each data point represents the average of three independent experiments. Error bars represent sd. (C) RNA synthesis of POLRMT, POLRMT^E423P^ (500 fmol) and T7 RNA polymerase (0.5 units) on M13mp18 ssDNA template. RNA products were separated on a denaturing polyacrylamide gel (10%). Molecular size markers are shown on the left. (D) Kinetics of Biotin-16-UTP incorporation in the *in vitro* transcription assay using POLRMT, POLRMT^E423P^ (500 fmol) or T7 RNA polymerase (0.5 units) on ssDNA template. Each time point represents the average of three independent experiments. Error bars represent sd. A line of linear regression was plotted for each protein. (POLRMT, R^2^=0.98; POLRMT^E423P^, R^2^=0.84; T7 RNA polymerase, R^2^=0.96.) (E) The RNA-primed DNA synthesis assay was monitored in the presence of Biotin-11-dCTP for labeling of DNA products. Lane 1, POLRMT (500 fmol) and POL γ (300 fmol); Lane 2, POLRMT^E423P^ (500 fmol) and POL γ (300 fmol); Lane 3, Control experiment using POL γ (300 fmol) only. Reactions were incubated for 1 h at 37 °C and the products were analyzed on a denaturing polyacrylamide gel (10%). Each data point represents the average of three independent experiments. Error bars represent sd.

### Exoribonuclease activity of POLRMT is required for short RNAs synthesis to prime DNA replication

In *Drosophila*, besides POLRMT, there are 4 additional PPR proteins encoded by *CG4611*, *CG10302*, *CG4679* and *CG14786* (Figure S3B). Phylogenetic analysis revealed that the PPR domain of fly POLRMT was closely related to PPR domains of POLRMTs from other species rather than other fly PPRs (Figure S3B). We were intrigued whether the exoribonuclease activity is a special feature of POLRMT or a general property of all PPR proteins. We expressed and purified full length proteins of CG10302, CG4679, CG14786, CG4611 and PPR domain of human POLRMT (hPOLRMT-PPR), and assayed these recombinant proteins for potential ribonuclease activity. None of the other 4 fly PPR proteins showed any RNase activity (Figure S3D, S3F), whereas hPOLRMT-PPR degraded 5’-Biotin-labeled ssRNA in a pattern similar to that of fly POLRMT (Figure S3E, S3F). Therefore, the exoribonuclease activity seems specific to animal mitochondrial RNA polymerase and may play conserved roles in mitochondrial DNA or RNA metabolism.

The PPR deletion mutant of POLRMT seemed unstable as its expression level in *E. coli* was extremely low, and no protein was detected in transgenic flies expressing this deletion mutant. We thus conducted a systemic mutagenesis, seeking for point mutations that are stable, and affect the exoribonuclease activity specifically. Sequence alignment of PPR domains in POLRMTs from different species revealed 22 identical or highly conserved residues forming a hydrophobic core of the helix-turn-helix fold (Figure S3C). We found that POLRMT with a single amino acid residue substitution of Glu-423 to Pro in the PPR domain (POLRMT^E423P^) completely abolished the exoribonuclease activity (Figure 3A, 3B). POLRMT^E423P^ was stable, and fully retained the polymerase activity in an *in vitro* transcription assay (Figure 3C, 3D). Importantly, POLRMT^E423P^, similar to T7 RNA polymerase and the PPR deletion mutant, synthesized long RNA species only (Figure 3C), further substantiating that the exoribonuclease activity is necessary for POLRMT to produce short RNA species.

Using ssDNA as a template, human POLRMT, similar to fly POLRMT, preferentially produces short RNA species that can prime DNA synthesis in the presence of POL γ (Wanrooij et al., 2008). We next asked whether fly POLRMT can also prime DNA synthesis. In a reaction mix containing POL γ, M13 ssDNA template, NTPs and dNTPs, the addition of POLRMT led to effective incorporation of Biotin-11-dCTP into DNA fragments that were longer than the short RNA species produced by POLRMT (Figure 3E). However, the incorporation of dNTPs was much less in the presence of POLRMT^E423P^ compared to POLRMT, and no obvious size shift was observed (Figure 3E,). These results demonstrate that POLRMT synthesizes short RNA oligos, which can effectively prime DNA replication *in vitro*.

### Exoribonuclease activity of POLRMT is required for mtDNA replication

To further understand the function of POLRMT’s ribonuclease activity *in vivo*, we generated transgenes expressing *POLRMT* or *POLRMT^E423P^* under the control of the *UASz* promoter (DeLuca and Spradling, 2018). When ubiquitously expressed in *POLRMT^del^* flies activated by *Actin-gal4*, *POLRMT* restored the steady-state levels of mtDNA and mitochondrial transcripts (Figure 4A), and fully rescued the viability of mutant flies. In contrast, *POLRMT^E423P^* failed to alleviate any of these defects (Figure 4A), indicating that exoribonuclease activity is critical for mtDNA maintenance. Given that POLRMT^E423P^ retains polymerase activity, the lack of rescue on mtDNA transcription could be a secondary effect of mtDNA deficiency in mutant flies.

**Figure 4.**
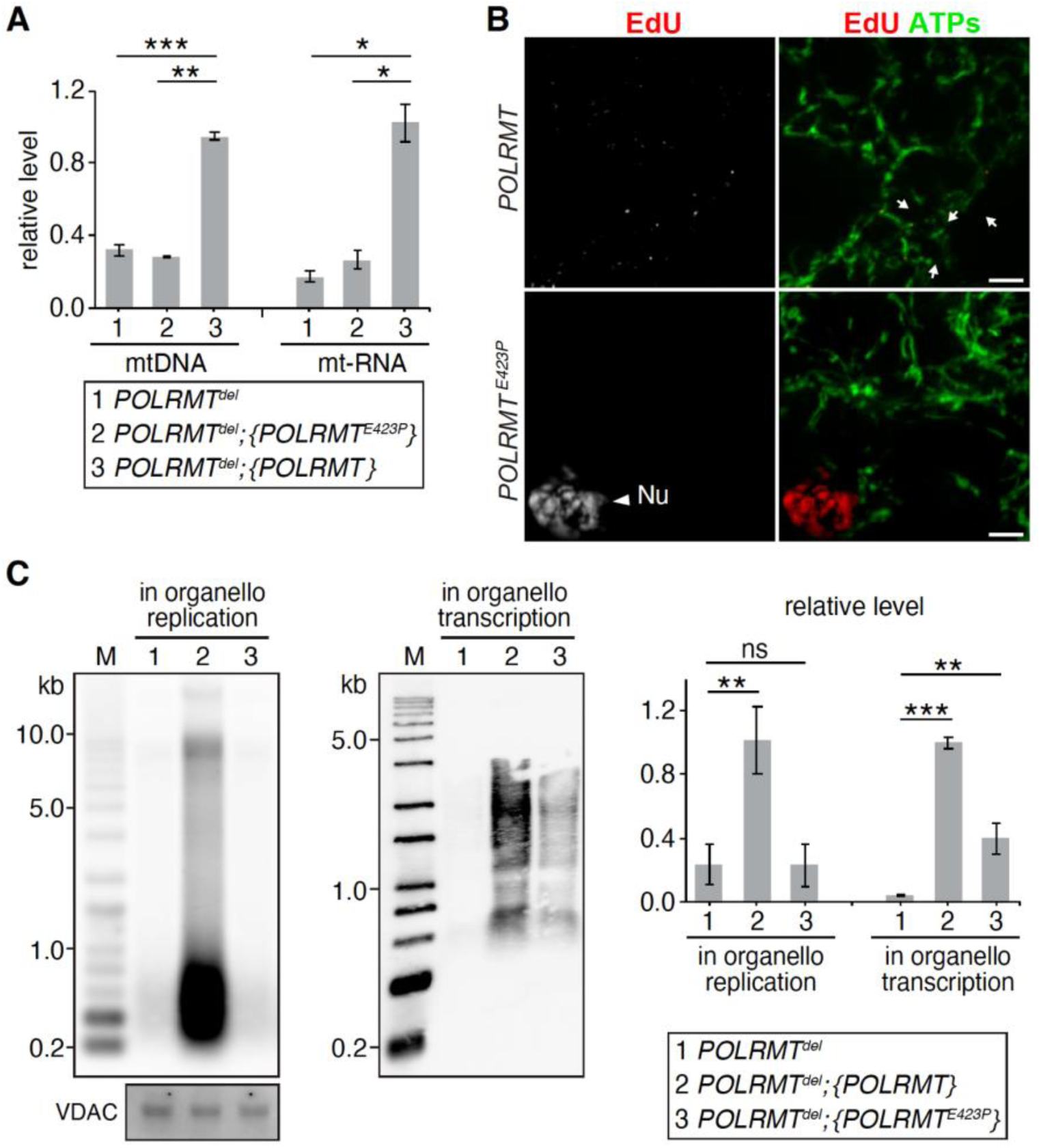
Exoribonuclease activity of POLRMT is required for *de novo* mtDNA synthesis. (A) Real-time PCR quantification of relative levels of mitochondrial DNAs and mRNAs in *POLRMT*^del^, *POLRMT*^del^ expressing POLRMT or POLRMTE^423P^ larvae. Each data point represents the average of three independent experiments. Error bars represent sd. *p<0.01; **p<0.001; ***p<0.0001. (B) Representative images of larva fat body from homozygous *POLRMT^del^* expressing full-length *POLRMT* and *POLRMT^E423P^* that were fed with EdU (Red) and co-stained for ATP synthase (ATPs; Green). Arrows indicate EdU incorporation in mitochondria; arrowhead indicates EdU incorporation in the nucleus (Nu) of *POLRMT^E423P^* fat body cells. Scale bars, 10 µm. (C) *In organello* mtDNA replication and transcription in mitochondria purified from *POLRMT^del^*, *POLRMT^del^* expressing POLRMT or POLRMT^E423P^ flies. Mitochondria isolated from third-instar larvae were incubated with Biotin-11-dCTP or Biotin-16-UTP to label *de novo* synthesized mitochondrial DNA and RNA respectively. Equal amount of mitochondrial preparation was loaded and validated by VDAC. Each data point represents the average of three independent experiments. Error bars represent sd. *p<0.01; **p<0.001; ***p<0.0001. Genotypes in (A) and (C): 1. *POLRMT^del^*, *w;POLRMT^del^/POLRMT^del^.* 2. *POLRMT^del^; {POLRMT^E423P^}, w;POLRMT^del^/POLRMT^del^;ac-Gal4>UAS-POLRMT^E423P^*. 3. *POLRMT^del^; {POLRMT}, w;POLRMT^del^/POLRMT^del^; ac-Gal4>UAS-POLRMT*. Genotypes in (B): *POLRMT, w;POLRMT^del^/POLRMT^del^; ac-Gal4>UAS-POLRMT. POLRMT^E423P^*, *w;POLRMT^del^/POLRMT^del^; ac-Gal4>UAS-POLRMT^E423P^*.

To determine whether the reduced steady-state levels of mtDNA and mitochondrial transcripts are caused by defective *de novo* synthesis, we carried out *in organello* mtDNA replication and transcription assays. Mitochondria isolated from *POLRMT^del^* flies lacked *de novo* synthesis of either mtDNA or mitochondrial transcripts, both of which were restored by the expression of wild type *POLRMT* (Figure 4C). A group of low molecular weight DNA fragments around 500 bp that resembles mammalian 7S DNA was also emerged from the *in organello* mtDNA replication assay (Figure 4C), suggesting a potentially conserved mechanism of mtDNA replication between mammals and flies. The expression of *POLRMT^E423P^* partially restored *in organello* synthesis of mitochondrial transcripts, but not mtDNA replication (Figure 4C), which is consistent with the notion that E423P mutation does not interfere with the polymerase activity. We also performed EdU incorporation assay to directly assess *de novo* mtDNA replication in larva fat body. Newly synthesized mtDNA marked as EdU puncta in mitochondria were detected in *POLRMT^del^* flies expressing *POLRMT, but not POLRMT^E423P^* (Figure 4B). Together, these results confirm that the ribonuclease activity of POLRMT is indeed essential for the *de novo* synthesis of mtDNA.

### Overexpression of exoribonuclease deficient POLRMT causes neuromuscular defects and premature aging

POLRMT^E423P^ appeared to be dominant negative, as its overexpression in wild type background caused lethality at larval stage. To bypass this development defect and to further explore its impact on mitochondria and whole animal physiology, we applied “GeneSwitch” inducible Gal4 system (Roman et al., 2001), to express POLRMT^E423P^ in adult flies only (Figure S4C). As the control, flies overexpressing wild type POLRMT were largely normal (Figure 5A-C). Overexpression of POLRMT^E423P^ led to a gradual decline of mobility (Figure 5A), and markedly reduced life span (Figure 5B). One week after the induction of POLRMT^E423P^, female flies become semi-sterile. Their eggs had reduced mtDNA level and most of them failed to hatch (Figure 5C). These defects resemble phenotypes of *mdi* mutant that has impaired mtDNA replication in ovaries (Zhang et al., 2016), further substantiating a role of exoribonuclease activity in mtDNA replication. Two weeks after the induction, POLRMT^E423P^ expressing flies showed reduced respiration (Figure 5D). The cytochrome C oxidase activity was also markedly reduced in POLRMT^E423P^ flies (Figure 5E), while the succinate dehydrogenase that are solely encoded by nuclear genome was normal (Figure 5E), indicating that mitochondrial defects are pertained to mtDNA-encoded genes specifically.

**Figure 5.**
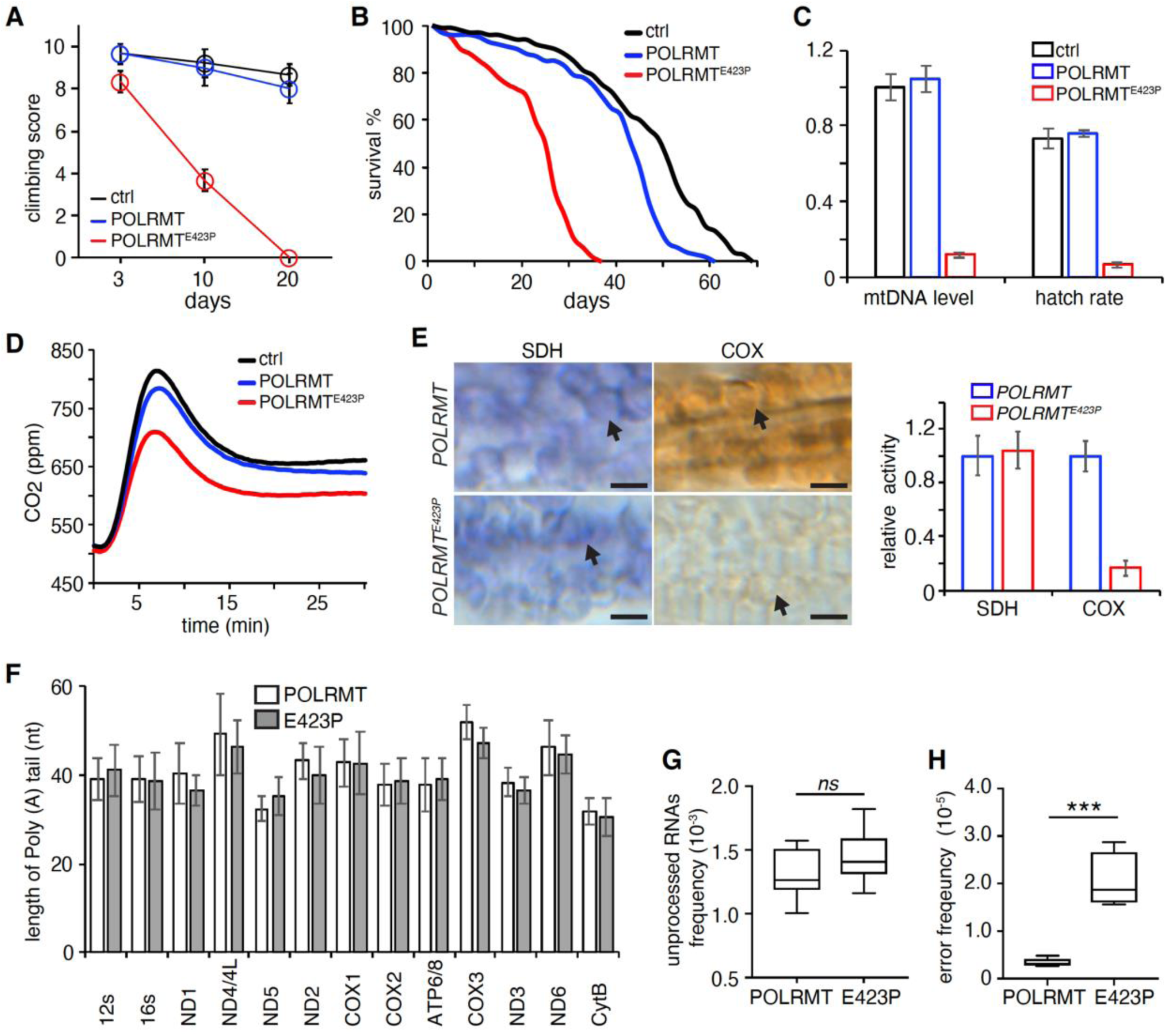
Expression of *POLRMT^E423P^* transgene in adult flies impairs mitochondrial activity and animal fitness. (A) Climbing assay of flies with indicated genotypes at different ages. Each data point represents the average of 10 individual flies (means ± SD, n = 3). (B) Lifespan of flies with indicated genotypes at 25 °C. n=200 flies for each genotype. (C) Hatching rate and mtDNA level of eggs laid by females of different genotypes (means ± SD, n = 3). (D) Representative graphs of CO_2_ production by male flies of different genotypes. At least three independent experiments were performed. (E) Representative images and quantification of adult muscles stained for SDH or COX activity. SDH or COX activity is normalized to that in the POLRMT overexpression muscles. n = 10 for each genotype. Arrows indicate mitochondria. Scale bars, 5 µm. (F) Quantification of the polyadenylation status of mRNAs in mitochondria. Error bars represent sd. (G) Frequency of the unprocessed mitochondrial transcripts. Error bars represent sd. p>0.05. (H) Error rate of mitochondrial transcripts assayed by circle-RNA sequencing. The error frequency of mitochondrial transcripts in POLRMT^E423P^ flies was markedly increased. Error bars represent sd. **p<0.01. Genotypes in (A) to (E): Ctrl, *w;;ac-Gal4:PR*. POLRMT*, w;;ac-Gal4:PR>UAS-POLRMT.* POLRMT^E423P^*, w;;ac-Gal4:PR>UAS-POLRMT^E423P^*. Genotypes in (F) to (H): POLRMT, *w;;ac-Gal4:PR>UAS-POLRMT.* E423P, *w;;ac-Gal4:PR>UAS-POLRMT^E423P^*.

We were puzzled by the severe neuromuscular phenotypes of POLRMT^E423P^ overexpressing flies, as mtDNA replication events were rare in adult muscles compared to mitotic tissues (Figure S4A). Consistent with this notion, neither the number nor the intensity of TFAM-GFP puncta was affected in POLRMT^E423P^ flies (Figure S4B, S4D-F). Therefore, the exoribonuclease activity of POLRMT likely regulates some processes that are not directly related to mtDNA replication in adult muscles.

### Exoribonuclease activity of POLRMT contributes to mtDNA transcription proofreading

Mitochondrial RNAs are initially transcribed as long polycistronic precursors, and then processed into individual mRNAs, rRNAs and tRNAs (Ojala et al., 1981). Other PPR proteins have been shown to involve in mitochondrial RNA processing, editing and stability control (Bratic et al., 2011; Xu et al., 2004). We hence carried out additional analyses to examine potential impact of POLRMT^E423P^ on these processes. We performed single molecular FISH against mitochondrial RNAs and found their steady-state levels were comparable between POLRMT^E423P^ expressing flies and the control (Figure S5A-C). We also cloned and sequenced all 13 mRNAs and 2 rRNAs encoded on mtDNA. They all had corrected 3’ends and proper lengths of Poly A tail in POLRMT^E423P^ flies (Figure 5F). We then conducted paired-end RNA sequencing to evaluate mitochondrial RNA processing by quantifying the frequency of a single read spanning two adjacent genes. In control flies, 1,416 reads out of total 1,081,246 reads that aligned to fly mitochondrial genome were mapped to two adjacent genes, indicating a frequency of unprocessed preRNAs as 0.13%, which is similar to that in POLRMT^E423P^ flies (0.14%, 809 out of total 557597 reads, Figure 5G, Table S1). Therefore, the ribonuclease activity of POLRMT does not contribute to pre-RNA processing. Very few tRNA-derived sequences were emerged in the paired-end RNA sequencing, due to a lack of specific procedure to recover tRNAs during library preparation (Dard-Dascot et al., 2018). We thus performed Hydro-tRNAseq to evaluate the expression of mitochondrial tRNAs. Among 22 tRNAs encoded by mitochondrial genome, tRNA^Gln^, tRNA^his^ and tRNA^Pro^ had less than two-fold difference (Table S2), while the remaining 19 showed no difference between control and POLRMT^E423P^ flies (Table S2). It is unlikely this minor change of tRNA expression causes severe mitochondrial dysfunctions in POLRMT^E423P^ flies.

An evolutionary conserved transcription elongation factor, TFIIS stimulates the intrinsic exonucleolytic activity of RNA Polymerase II, which contributes to the transcription proofreading of nuclear genome (Kettenberger et al., 2003). We were intrigued whether the exoribonuclease activity of POLRMT might have a similar role in mitochondria. Considering that the actual errors on RNAs could be masked by reverse transcription errors during library preparation of regular RNAseq procedures, we adopted a “circle-sequencing method”, which sequences a cDNA molecule consists of a tandem repeat of a single RNA fragment (Acevedo and Andino, 2014). We analyzed more than 100 million reads from control and POLRMT^E423^ flies and obtained 500,000-fold coverage of the mitochondrial genome. In control flies, the error rate of mitochondrial transcripts was about 3.5×10^−6^ (Figure 5H), which is comparable to the error rate on nuclear transcripts at 4.0×10^−6^ (Gout et al., 2017). About 60% of transcription errors were base substitutions and the remaining were frameshifts. We did not notice any hot spots on mitochondrial genome for transcription errors, as the error frequency within a given RNA is proportional to its steady-state level (Figure S6A, 6B). Of primary significance, the error frequency of mitochondrial transcripts in POLRMT^E423P^ flies was 2.0×10^−5^, about 6 folds higher than that of control (Figure 5H), suggesting that the exoribonuclease activity of POLRMT contributes to mtDNA transcription proofreading.

## Discussion

Pentatricopeptide repeat proteins belong to a large protein family that predominantly localize to either chloroplasts or mitochondria, two organelles having their own genomes (Rovira and Smith, 2019). Different from the nuclear genome, animal mitochondrial genomes lack regulatory sequences, and are transcribed as polycistronic RNA precursors (Anderson et al., 1981). Post-transcriptional mechanisms are not only essential to produce mature RNA species, but also play important roles in regulating organellar gene expression. PPR proteins have been predicted to bind to RNAs, and suggested to involve in RNA processing, editing and translational regulations in organelles (Lightowlers and Chrzanowska-Lightowlers, 2008). Nonetheless, the biochemical basis of PPR proteins in RNA metabolism is largely unknown. We, for the first time, demonstrate that a PPR protein, the *Drosophila* mitochondrial RNA polymerase possesses a 3’-5’ exoribonuclease activity. We found that PPR’s ribonuclease activity was insensitive to broad spectrum RNase inhibitor but required divalent cations, which are dispensable for common RNase contaminations (Fersht, 1977). Importantly, the degradation products of PPR protein had 3’-hydroxyl group, whereas the destructive RNases in contamination often generate 3’-phosphate group (D’Alessio and Riordan, 1997). We hence conclude that the detected ribonuclease activity of recombinant PPR protein is not due to RNase contamination, despite its prevalence in routine molecular biology experiments.

We found steady-state levels of both mtDNA and mitochondrial transcripts were significantly reduced in *POLRMT^del^* flies, similar to phenotypes of *POLRMT* knockout mice (Kühl et al., 2016). Importantly, the *de novo* mtDNA replication was completely abolished, while mtDNA nucleoids were still present in *POLRMT^del^* flies, demonstrating a direct role of POLRMT in mtDNA synthesis. Our results are in line with previous studies in mammals and support the idea that POLRMT is the mitochondrial primase. Similar to mammalian POLRMTs (Wanrooij et al., 2008; Kühl et al., 2016), recombinant *Drosophila* POLRMT also produced short RNA species. The deletion of PPR domain or the E432P point mutation generated much longer transcripts, indicating that the 3’-5’ exoribonuclease activity is necessary for the generation of short RNA oligos to prime mtDNA replication.

The point mutation, POLRMT^E423P^ completely abolished the exoribonuclease activity, but retained the polymerase activity. Overexpression of POLRMT^E423P^ led to developmental arrest at 3^rd^ instar larva and markedly inhibited *de novo* mtDNA replication, further supporting a critical role of exoribonuclease activity in mtDNA replication. When overexpressed in adult flies specifically, POLRMT^E423P^ caused severely neuromuscular defects that are typical features of mitochondrial disruptions. However, these phenotypes cannot be attributed to PPR’s role in mtDNA replication, as mtDNA is not actively replicated in post-mitotic tissues of adult flies. We found that the steady-state level and the processing of mature mitochondrial mRNAs and rRNAs were not affected in POLRMT^E423P^ overexpression flies. We only detected a noticeable, and consistent increase of RNA misincorporations in POLRMT^E423P^ flies compared to that in control, suggesting that the 3’-5’ exoribonuclease activity of POLRMT may contribute to mitochondrial transcription proofreading.

To ensure the survival and continuity of life, the flow of hereditary information from DNA to RNA and then to protein demands remarkable accuracy. Even though transcription errors on RNAs are transient, they are much more frequent compared to DNA replication errors (Kirkwood, 2012). One erroneous mRNA molecule can be translated to multiple copies of aberrant proteins that would profoundly affect cellular fitness. Nuclear genome relies on both transcriptional proofreading and mRNA surveillance mechanisms to ensure the fidelity of mRNAs (Moraes, 2010; Thomas et al., 1998). To our knowledge, the potential proofreading mediated by POLRMT’s exoribonuclease activity represents the only mechanism safeguarding the information flow from DNA to RNA in mitochondria. Proofreading of mtDNA replication is carried out by the intrinsic 3’-5’ exonuclease activity of polymerase γ (Bratic et al., 2015). The loss of such activity drastically elevates mtDNA mutations frequency and causes premature aging phenotypes or developmental arrest in engineered mice and flies respectively (Bratic et al., 2015; Trifunovic et al., 2004). In fact, constant overexpression of POLRMT^E423P^ led to developmental arrest at 3^rd^ instar larva, while restricted expression in adulthood caused severe premature aging phenotypes, both of which resemble that of animals with elevated mtDNA mutations.

While PPRs share little homology on primary sequence, phylogenetic analyses reveal that PPR domains of animal mitochondrial RNA polymerase are closely related to each other (Figure S3B). Noteworthy, the PPR domain of human POLRMT also contained 3’-5’ exoribonuclease activity. We notice that the number of PPR proteins is positively correlated with the size of organelle genome in a given organism (Figure S3A). In various lineages of land plants, which have much larger and more complex organellar genomes, PPR protein family has expanded to over 400 members (Fauron et al., 2004; Fujii and Small, 2011; Palmer, 1985). It is highly possible that some PPR proteins, besides animal mitochondrial RNA polymerases, might have ribonuclease activities as well. The whole picture of PPR domain and its ribonuclease activity in mitochondrial RNA metabolism awaits future investigations.

### Limitations

We demonstrate that the 3’-5’ exoribonuclease activity of PPR domain of POLRMT is necessary for the synthesize short RNA oligos to prime mtDNA replication. However, it is not clear how the ribonuclease activity stimulates the production of short RNA species. The PPR domain locates at the RNA exist channel of POLRMT-DNA transcription complex (Ringel et al., 2011). It might degrade newly synthesized RNA in a 3’-5’ direction, and trigger the release of POLRMT from DNA template. The RNA oligos with 3’ hydroxyl group therein can be utilized to prime DNA synthesis. Additionally, it remains a question what determines the switch between continuous transcription and mtDNA replication. In mammals, the mitochondrial transcription elongation factor, TEFM stimulates POLRMT polymerase activity, and encourages the continuous synthesis of long RNAs (Agaronyan et al., 2015). It is possible that TEFM might inhibit POLRMT ribonuclease activity to promote transcription, or other factors may act in an opposite way when mtDNA replication is demanded.

## STAR★METHODS

Detailed methods are provided in the online version of this paper and include the following:

- KEY RESOURCES TABLE
- CONTACT FOR REAGENT AND RESOURCE SHARING
- EXPERIMENTAL MODEL AND SUBJECT DETAILS
  - Fly genetics and husbandry
  - Cell culture
- METHODS DETAILS
  - Life span assay
  - Climbing assay
  - RNA/DNA isolation of quantitative real-time PCR
  - EdU incorporation
  - Immunostaining and single-molecule fluorescence in situ hybridization
  - Mitochondrial isolation
  - Protein expression and purification
  - *In vitro* transcription assay
  - *In vitro* RNA-primed DNA replication
  - *In organelle* replication and transcription assays
  - Ribonuclease activity assay
  - Poly(A) tail length assay
  - Electrophoretic mobility shift assay (EMSA)
  - SDH and COX activity assay
  - RNA sequencing and analyses
  - Confocal Microscopy
- QUANTIFICATION AND STATISTICAL ANALYSIS
- DATA AND SOFTWARE AVAILABILITY

## KEY RESOURCES TABLE

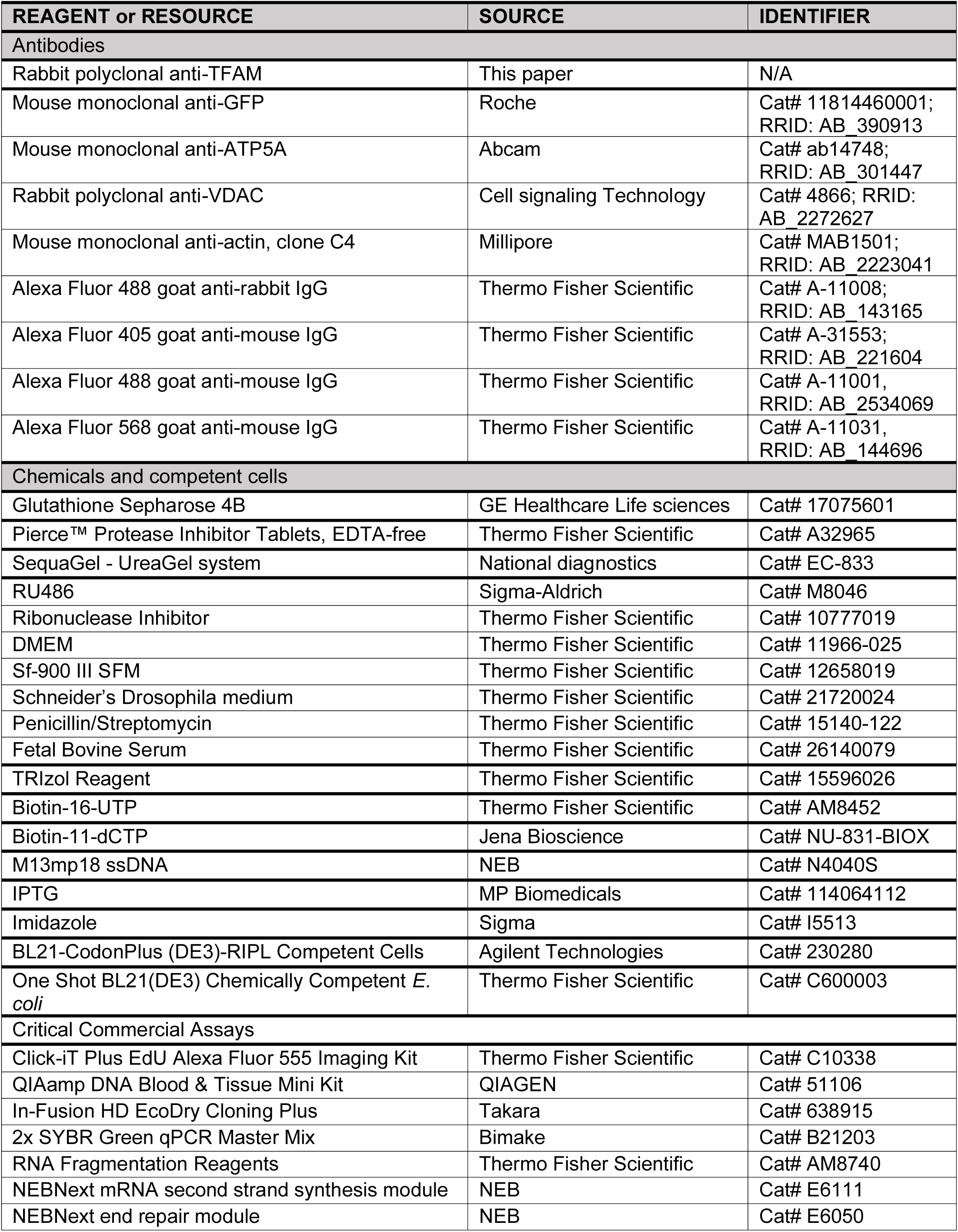
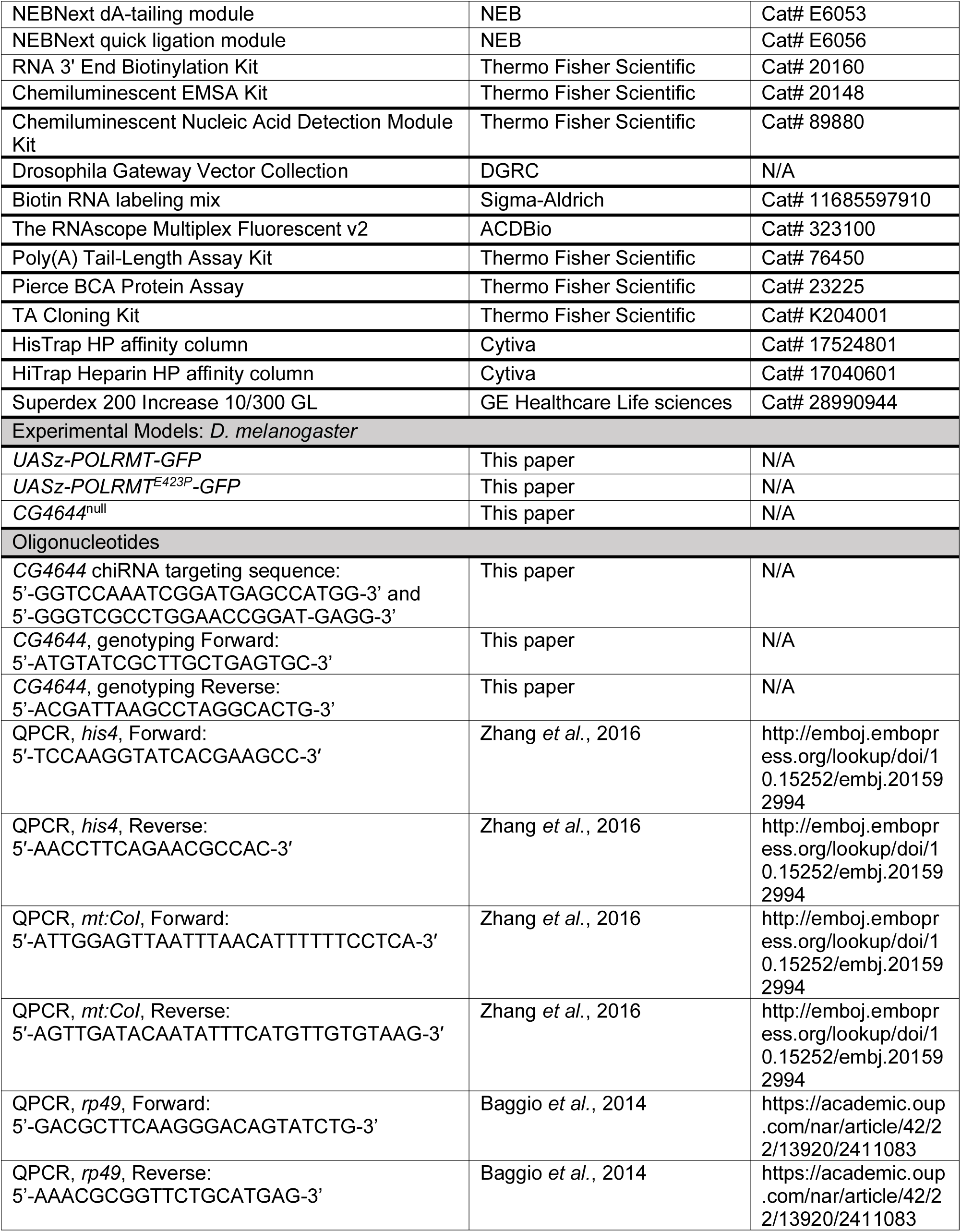
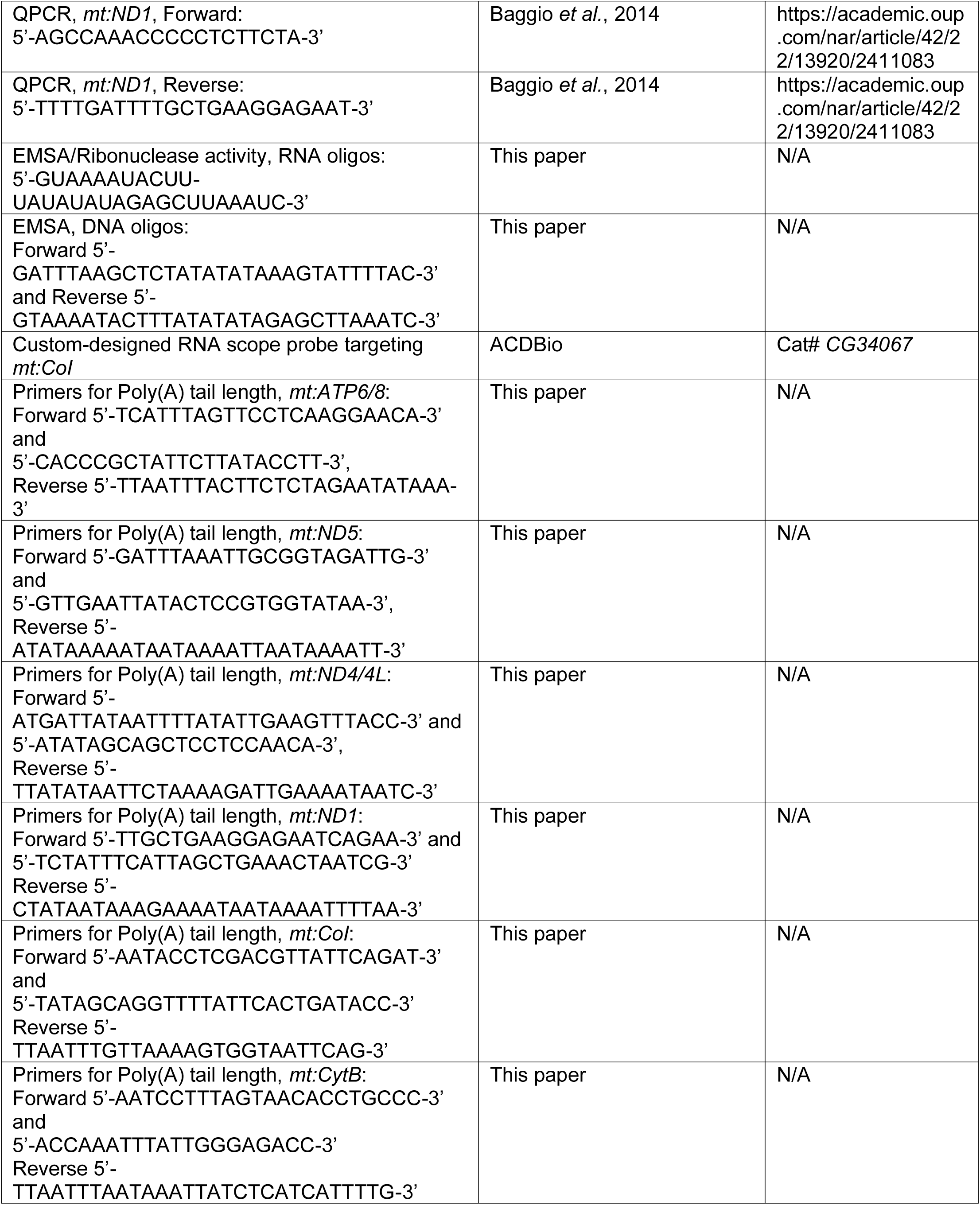
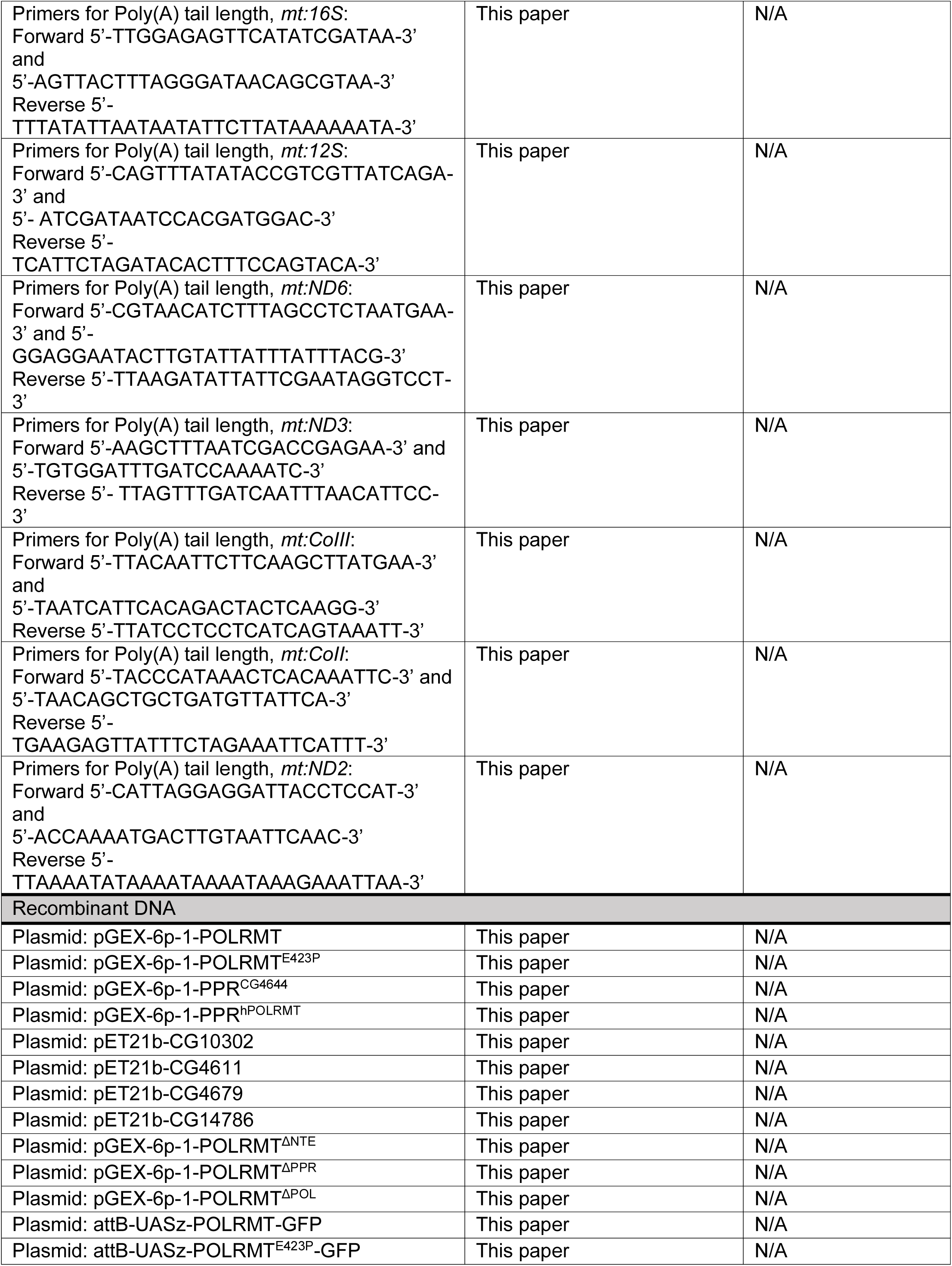
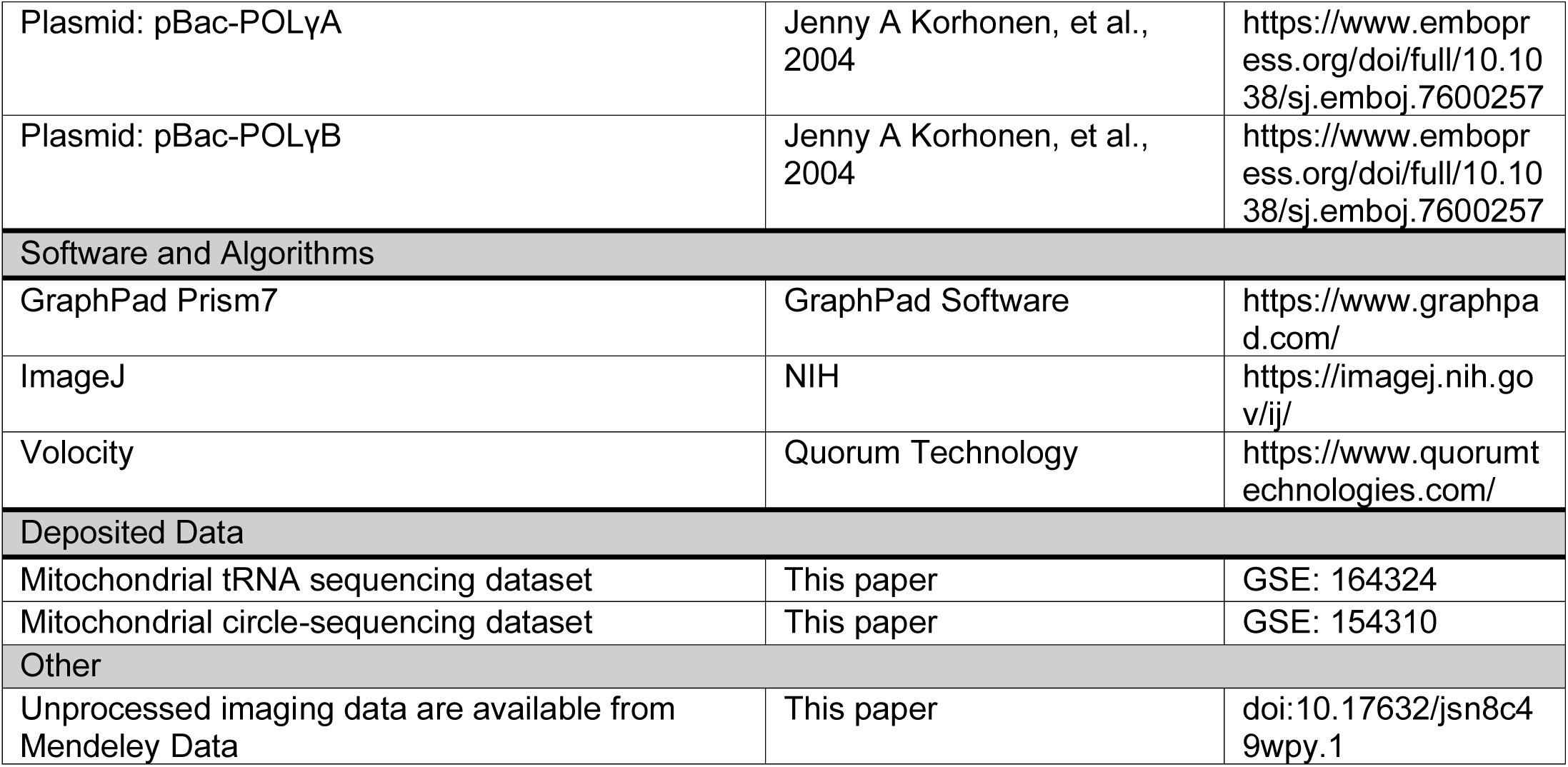

## RESOURCE AVAILABLITY

Further information and requests for resources and reagents should be directed to and will be fulfilled by the Lead Contact, Hong Xu.

## Materials Availability

Materials such as plasmids generated in this study are available upon request from the lead contact.

## Data and Code Availability

The RNA sequencing data generated in this study have been deposited at the Gene Expression Omnibus of NCBI with accession number (GEO: GSE154310 and GSE 164324). Unprocessed imaging data (microscopy as well as blots) are available at Mendeley Data: doi:10.17632/jsn8c49wpy.1.

## EXPERIMENTAL MODEL AND SUBJECT DETAILS

### *Drosophila* genetics and husbandry

Flies were maintained on standard cornmeal-yeast medium at 25 °C. BL9431(Gal4:PR) line was obtained from Bloomington Stock Center. Transgenic flies, *UASz-POLRMT-gfp*, *UASz-POLRMT^E423P^-gfp* were injected by the BestGene embryonic injection service. Inducible transgene expression was activated by transferring adult flies to fly food containing RU486 (Sigma). The final effective concentration of RU486 is 500 µM. The deletion mutant of *Drosophila* mitochondrial RNA polymerase encoded by *CG4644* locus was generated by CRISPR/Cas-9-mediated recombination. Donor plasmid and chiRNAs (5’GGTCCAAATCGGATGAGCCATGG3’ and 5’GGGTCGCCTGGAACCGGATGAG3’) were injected into the embryos of PBac{y[+mDint2] = vas-Cas9} VK00027 flies by the BestGene embryonic injection service. G1 adults were screened for deletion events by PCR (5’ATGTATCGCTTGCTGAGTGC3’ and 5’ACGATTAAGCCTAGGCACTG3’). One line of 1.5 kb deletion that removes the majority of *CG4644* genomic region including an exon encoding the polymerase domain was identified and named as *POLRMT^del^*. *POLRMT^del^* was backcrossed with *w^1118^* for 5 generations to clean up the genetic background. *w^1118^* was used as the wild type control, or otherwise indicated.

### Cell culture

*Drosophila* S2 cells were cultured in Schneider’s *Drosophila* medium (Invitrogen) supplemented with 10% FBS and 1% penicillin-streptomycin at 25 °C. HeLa cells were maintained in Dulbecco’s modified Eagle medium (GIBCO) supplemented with 10% FBS and 1% penicillin-streptomycin at 37 °C with 5% CO_2_. Sf9 insect cells were cultured in sf-900 III SFM (Invitrogen) at 27 °C with shaking at 140 rpm. Lipofectamine 3000 reagent (Thermo Fisher Scientific) was used for plasmid transfection.

## METHOD DETAILS

### Lifespan assay

Newly-eclosed male flies were collected and maintained at 25 °C. A group of 20 flies with the same genotype were placed into an individual vial and transferred to a new vial every other day. At least 10 replicates were set up for each genotype. The number of dead flies was recorded daily, and the survivorship curve was displayed by the percentage of total flies surviving.

### Climbing assay

In the climbing test, 10 male flies were placed into a transparent glass tube. 3 replicate tubes, containing 10 flies each were tapped on bench simultaneously. The flies were allowed to climb up a 15-cm predefined height and the time for flies to climb up was 60 s. The number of flies in each tube was counted. The climbing score was calculated by averaged number of flies in replicate vials. All tests were performed at the same time of a day to minimize circadian rhythm effects.

### RNA/DNA isolation and quantitative real-time PCR

Total RNA from *Drosophila* tissues was isolated using TRIzol reagent (Invitrogen). The isolated RNA was treated with DNase I (Invitrogen) at room temperature for 15 min. The treated RNA was further cleaned up using a RNeasy Mini Kit (Qiagen). A Superscript First-strand Synthesis kit (Invitrogen) was used for cDNA synthesis. One-step quantitative real-time PCR was performed on a Lightcycler 480 II PCR system (Roche) by using SYBR Green qPCR SuperMix (Bio-rad). To quantify mtDNA level, total DNA of third-instar larvae was isolated using DNeasy Blood & Tissue Kit (Qiagen). qPCR was performed as described previously (Zhang et al., 2015).

### EdU incorporation

To assay the mtDNA replication in adult tissues, EdU staining was performed using the EdU feeding assay with some modifications (Micchelli and Perrimon, 2006). Flies were reared on standard fly media with 0.25 mM EdU at 25 °C. The tissues were dissected and assayed with Click-iT Plus EdU Imaging Kit (Thermo Fisher Scientific).

### Immunostaining and single-molecule fluorescence in situ hybridization assay (sm-FISH)

Sm-FISH was carried out as described previously (Chen et al., 2020). Sm-FISH RNA probe for *mt:CoI* was synthesized by ACDBio and the whole-mount in situ hybridization procedure was performed on adult indirect flight muscles as described previously (Pharris et al., 2017). The tissue was co-stained with an Anti-ATP5A antibody (Abcam, 15H4C4, 1:1000) in the dark at 4 °C overnight. The secondary antibody, Alexa Fluor 568 goat anti-mouse IgG (Invitrogen) was used at 1:200 in blocking solution for 2 hr at room temperature.

### Mitochondria isolation

Mitochondria isolation was performed as previously described with modifications (Chen et al., 2014). Adult flies were collected on ice and gently homogenized by plastic microtube pestles in ice-cold isolation buffer. The homogenate was then transferred to a 1 mL syringe and forced through the cotton filter. The collected fluid was centrifuged at 500 × *g* for 3 min at 4 °C. The resulted supernatant was centrifuged at 9000 × *g* for 10 min. The pellet was collected and washed twice with the isolation buffer.

### Protein expression and purification

The GST-tagged *Drosophila* PPR recombinant proteins including PPR, PPR^E423P^, POLRMT, POLRMT^E423P^, ΔNTE, ΔPPR and ΔPOL were expressed in *E. coli* BL21(DE3) cells (Invitrogen). Crude cell lysate was treated with 0.6% (w/v, pH 8.0) Polyethyleneimine (MP Biomedicals) to precipitate the nucleic acids. The supernatant was then loaded on Glutathione Sepharose 4B column (GE Healthcare) according to the manufacturer’s protocol. The affinity-purified proteins were further passed through an ion exchange column and a size-exclusion column employed on ÄKTA pure protein purification system.

Human POL γ A and POL γ B were expressed in *Spodoptera frugiperda* (Sf9) cells and purified as described previously (Korhonen et al., 2004). Sf9 cells were co-infected with recombinant baculoviruses encoding POL γ subunits A and B (gifted from Dr. Maria Falkenberg, Department of Medical Nutrition, Karolinska Institute, Novum, Huddinge, Sweden). Whole-cell extract was loaded on HisTrap column and HiTrap heparin column (GE Healthcare). The POL γ A subunit was eluted as a heterodimeric complex with POL γ B and the holoenzyme containing both subunits is referred as POL γ.

To generate CG4611, CG4679, CG10302 and CG14786 recombinant proteins, the coding regions (without targeting peptides) were amplified by PCR from *Drosophila* cDNA, cloned into the expression vector pET21b, and expressed using BL21-CodonPlus (DE3)-RIPL cells (Agilent). All proteins were first purified using affinity chromatography on HisTrap column followed by HiTrap heparin purification in 250-1500 mM gradient of NaCl (GE Healthcare). Protein concentration was determined by the Pierce BCA reagents (Thermo Fisher Scientific), with BSA as the standard.

### *In vitro* transcription assay

The *in vitro* transcription assay on ssDNA template was carried out as described previously (Wanrooij et al., 2008). The 20 µl reaction mix containing 80 ng of single-stranded M13mp18 DNA (New England Biolabs), 10 mM Tris·HCl (pH 8.0), 100 µg/ml BSA, 20 mM MgCl_2_, 1 mM DTT, 1 mM each ATP, CTP and GTP, 0.65 mM UTP, 0.35 mM Biotin-16-UTP (Sigma), 10 units of RNase inhibitor (Thermo Fisher Scientific), with the indicated amount of T7 RNA polymerase (Ambion) or 500 fmol of purified recombinant proteins was incubated at 32 °C for 30 min. The RNA products were extracted twice with Phenol: Chloroform: Isoamyl Alcohol 25:24:1 (Sigma) and precipitated with 2.5 - 3.0 volumes of ethanol for 1 h to overnight at −20 °C. The samples were dissolved in RNA sample loading buffer (Sigma), denatured for 5 min at 75 °C, and analyzed on a denaturing polyacrylamide gel. The products were detected by the Chemiluminescent Nucleic Acid Detection Module (Thermo Fisher Scientific). To semi-quantify the level of *in vitro* transcription, the total densitometry of Biotin-16-UTP chemiluminescence in each lane was measured and normalized to that of 0.1 pmol 5’-Biotin-ssRNA, which was loaded on the same gel and hence used as the internal standard. The relative level of Biotin-16-UTP in each reaction was calculated and plotted.

### *In vitro* RNA-primed DNA replication

The *in vitro* RNA-primed DNA replication assay was carried out using M13mp18 ssDNA as the template and in the presence of Biotin-11-dCTP (Jena Bioscience). Reactions (20 µl) contained 35 fmol M13mp18 ssDNA, 10.5 mM MgCl_2_, 25 mM Tris-HCl (pH 8.0), 1 mM DTT, 100 µg/ml BSA, 400 µM ATP, 150 µM GTP, UTP and CTP, 100 µM dATP, dGTP and dTTP, 60 µM dCTP, and 40 µM Biotin-11-dCTP, 300 fmol POL γ and 500 fmol POLRMT or 500 fmol POLRMT^E423P^. After incubation for 1 h at 37 °C the DNA products were analyzed by electrophoresis on a denaturing polyacrylamide gel (10%) and detected by chemiluminescent detection (Thermo Fisher Scientific). To semi-quantify the level of dCTP incorporation, the total densitometry of Biotin-dCTP chemiluminescence in each lane was measured and normalized to the densitometry of 0.1 pmol 5’-Biotin-ssRNA that was loaded on the same gel and hence used as the internal standard. The relative level of Biotin-dCTP in each reaction was calculated and plotted.

### *In organello* replication and transcription assays

*In organello* replication and transcription assays were performed on mitochondria isolated from third-instar larvae as previously described (Enríquez et al., 1996; Gensler et al., 2001). For each sample, 1 mg of purified mitochondria was suspended in 500 µl of incubation buffer (75mM sorbitol, 25 mM sucrose, 2.5 mM malate, 10 mM glutamate, 10 mM K_2_HPO_4_, 100 mM KCl, 5 mM MgCl_2_, 50 µM EDTA, 1 mM ADP, 10 mM Tris-HCl (pH 7.4), and 1% (w/v) fatty acid-free BSA), and incubated with Biotin-11-dCTP/Biotin-16-UTP for 2 h at 37 °C. After labeling, mitochondria were collected at 13,000 g for 1 min and washed three times with 10 mM Tris-HCl (pH 6.8), 0.15 mM MgCl_2_ and 10% glycerol. For each *in organello* replication assay, the pellet was lysed with 2% SDS and total mtDNA was extracted by phenol/chloroform/isoamyl alcohol (25:24:1) (Sigma). After precipitation the labeled DNA were denatured at 95 °C for 15 min and separated by 0.8% agarose gel electrophoresis. For each *in organello* transcription assay, total mitochondrial RNA was isolated by TRIzol Reagent (Invitrogen) and separated on 1.2% agarose gel. The labeled nucleic acids signals were detected by chemiluminescent detection (Thermo Fisher Scientific). To semi-quantify the in organello mtDNA replication and transcription, the level the total densitometry of Biotin-dCTP chemiluminescence (in organello mtDNA replication) and Biotin-UTP chemiluminescence (in organello mtDNA transcription) in each lane was measured and normalized to the densitometry of 0.1 pmol 5’-Biotin-ssRNA that was loaded on the same gel and hence used as the internal standard. The relative level of Biotin-dCTP or Biotin-UTP in each reaction was calculated and plotted.

### Ribonuclease activity assay

Each reaction contained 0.5 pmol of Biotin-labeled 30mer RNA oligos (5’-GUAAAAUACUU-UAUAUAUAGAGCUUAAAUC-3’), synthesized by IDT. The reaction mixture (10 µl) was consisted of 10 mM HEPES (pH 7.3), 20 mM KCl, 1 mM MgCl_2_ and 1 mM DTT. The ribonuclease assay was initiated by adding 0.5 µM of purified protein to the reaction, incubated at 32 °C for indicated times, and terminated by adding RNA loading buffer (Sigma). The reaction mix was denatured for 5 min at 75 °C, resolved on a denaturing polyacrylamide gel and detected using the Chemiluminescent Nucleic Acid Detection Module (Thermo Fisher Scientific). The Biotinylated DNA markers were synthesized by IDT.

### Poly(A) tail length assay

The poly(A) tail lengths of each mitochondrial transcripts were examined using Poly(A) Tail-Length Assay Kit (Thermo Fisher Scientific). Total RNA was extracted from isolated mitochondria and added a number of guanosine and inosine residues to the 3’-ends. The tailed-RNAs were converted to cDNA using the newly added G/I tails. The gene-specific forward primer and the universal reverse primer were used for PCR amplification. PCR products were then cloned using the TOPO TA Cloning Kit for sequencing (Thermo Fisher Scientific).

### Electrophoretic mobility shift assay (EMSA)

EMSA was performed using LightShift Chemiluminescent EMSA Kit. Reaction mix consisted of 50 nM biotin-labeled probes and PPR protein at different concentrations (0-1 µM) was incubated 30 min at room temperature in the binding buffer. After incubation, each sample was mixed with equal volume of loading buffer and analyzed on a DNA retardation gel. DNA or RNA probe was detected in gel using Chemiluminescent Nucleic Acid Detection Module (Thermo Fisher Scientific).

### SDH and COX activity assay

For SDH and COX activity staining, experiment was conducted as previously described with minor modifications (Wang et al., 2019). Briefly, adult muscles were dissected in PBS and resuspended in either SDH staining buffer (0.4 mM phenazine methosulfate, 0.5 mM nitro blue tetrazolium, 42 mM succinic acid, 60 µM rotenone, 4 mM antimycin A and 2 mM KCN) for 10 min or COX staining buffer (4 mM 3,3’-diaminobenzidine, 2 µg/ml catalase, 200 µM cytochrome c, 84 mM malonate, 60 µM rotenone and 4 mM antimycin A) for 30 min at room temperature. Samples were washed three times with 50 mM phosphate buffer (pH7.4), fixed with 4% paraformaldehyde for 15 min and finally immersed in 90% glycerol for microscopy. Bright images were captured by Zeiss microscope. Quantification of relative activities in COX and SDH assay was carried out in ImageJ. Color images were first converted into gray scale images and ‘Color Threshold’ was used. Densitometry of selected mitochondrial area was measured, and the non-selected area was considered as background.

### RNA sequencing and analyses

RNA sequencing and tRNA sequencing were performed with isolated mitochondria of adult flies. Total RNA was extracted from isolated mitochondria by TRIzol Reagent (Invitrogen). For unprocessed RNA frequency, polyadenylated mitochondrial mRNAs were captured by the DNA Sequencing and Genomics Core, NHLBI, NIH. Deep sequencing of the isolated mitochondria was performed on the Hiseq 3000 (Illumina) according to the Illumina RNA-Seq protocol. Mapping and alignment were performed using Trim Galore! (v0.3.7) and BowTie. Uniquely mapped paired-end reads against the reference genome were used for the following analyses. Briefly, to calculate unprocessed polycistronic transcripts, SAMtools was applied to count the numbers of mapped reads spanning adjacent coding genes.

Error rates per position was determined by circle-sequencing method as previously described (Acevedo and Andino, 2014). Briefly, 1-5 µg of purified mitochondrial RNA was fragmented at 70 for 7.5 min. The fragmented RNA was then purified and circularized with T4 RNA ligase 1 (NEB). Tandem repeats were generated during first-strand synthesis at 42 °C for 30 min. Library cloning and amplification were performed by NEBNext mRNA second strand synthesis module (NEB), NEBNext end repair module (NEB), NEBNext dA-tailing module (NEB), and NEBNext quick ligation module (NEB) according to the manufacturer’s instructions. Libraries were sequenced for 300 cycles on Illumina MiSeq machine. RNA-seq reads generated by circle sequencing was analyzed by a pipeline previously developed (Gout et al., 2017).

The tRNA sequencing was performed by Arraystar Inc, Maryland. Mitochondrial tRNA were purified from total mitochondrial RNA and m1A&m3c demethylated, then partially hydrolyzed according to Hydro-tRNAseq method (Arimbasseri et al., 2015; Gogakos et al., 2017). Fragments of 10 - 40 nt for partially hydrolyzed and re-phosphorylated tRNAs were converted to small RNA sequencing libraries using NEBNext multiplex small RNA library prep set (NEB) for Illumina kit. The completed libraries were quantified by Agilent 2100 Bioanalyzer and diluted. Sequencing was performed on Illunima NextSeq 500 system. Raw sequencing reads that passed the Illumina chastity filter were used for the following analysis. Trimmed reads (with 5’, 3’ - adaptor bases removed) were aligned to mature-tRNA reference sequences allowing a maximum of 3 mismatches. The tRNA expression profiling was calculated based on uniquely mapped reads.

### Confocal microscopy

All images were captured on an UltraView Vox confocal imaging system (Perkin Elmer) and were processed with Volocity software. Quantification for the images was performed in ImageJ.

## QUANTIFICATION AND STATISTICAL ANALYSIS

All data were presented as the mean ± sd. P values were performed with Two-tailed Student’s t test built in GraphPad Prism 7. Statistical significance of difference was considered when p<0.05.

## SUPPLEMENTAL INFORMATION

Supplemental Information includes six figures and two tables.

## ACKNOWLEDGMENTS

We thank Dr. Rod Levine for his advice on protein purification and structural modeling; Dr. Maria Falkenberg for pBac-POLyA and pBac-POLyB plasmids; Dr. Jing-Wei Zhang for advices on riboproteins analyses; Bloomington *Drosophila* Stock Center for various fly stocks; the Developmental Studies Hybridoma Bank for antibodies; and Bestgene Inc for transgenic service. This work was supported by the Intramural Research Program of National Heart, Lung and Blood Institute.

## AUTHORS CONTRIBUTIONS

Y.L. and H.X. conceived the project and designed the experiments. Y.L., Z.C., Z.-H.W., K.D., J.T., D.-Y. L., and Y.-S. L., performed the experiments. Y.L., Z.-H.W., M.P., and H.X. analyzed the data. Y.L. and H.X. wrote the manuscript.

## DECLARATION OF INTERESTS

The authors declare no competing interests.

## Supplemental Information

**Figure S1.**
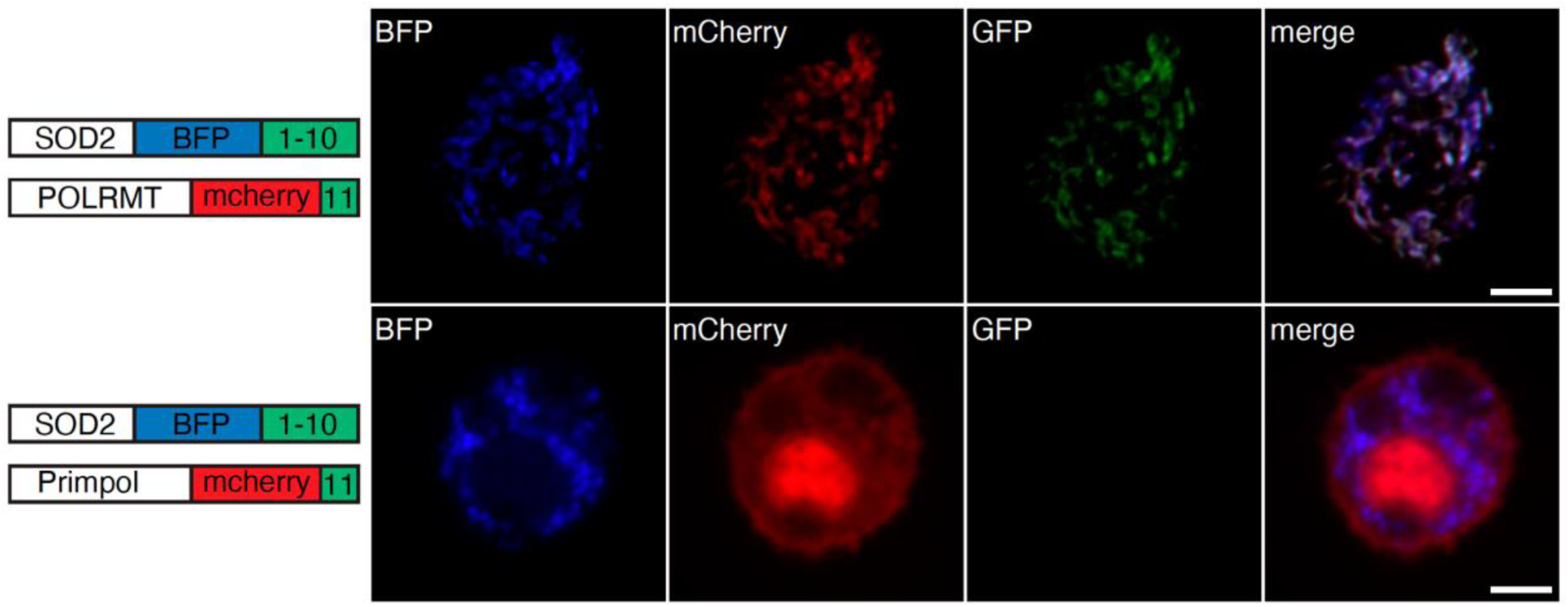
Human PrimPol is not localized in mitochondrial matrix. Left panel: Schematic representation of the constructs used in the GFP complementation assay. Right panel: Representative images of HeLa cell co-expressing SOD2-BFP-GFP1-10 (Blue) with human POLRMT-mCherry-GFP11 (Red); or human PrimPol-mCherry-GFP11 (Red). Note that GFP signal is evident and localizes to mitochondria in Hela cells expressing POLRMT-mCherry-GFP11 and SOD2-BFP-GFP1-10, indicating that both POLRMT and SOD2 localize to the mitochondrial matrix. The lack of GFP signal in Hela cells expressing SOD2-BFP-GFP1-10 and PrimPol-mCherry-GFP11, indicating that PrimPol does not localize to the mitochondrial matrix. Scale bars, 10 µm.

**Figure S2.**
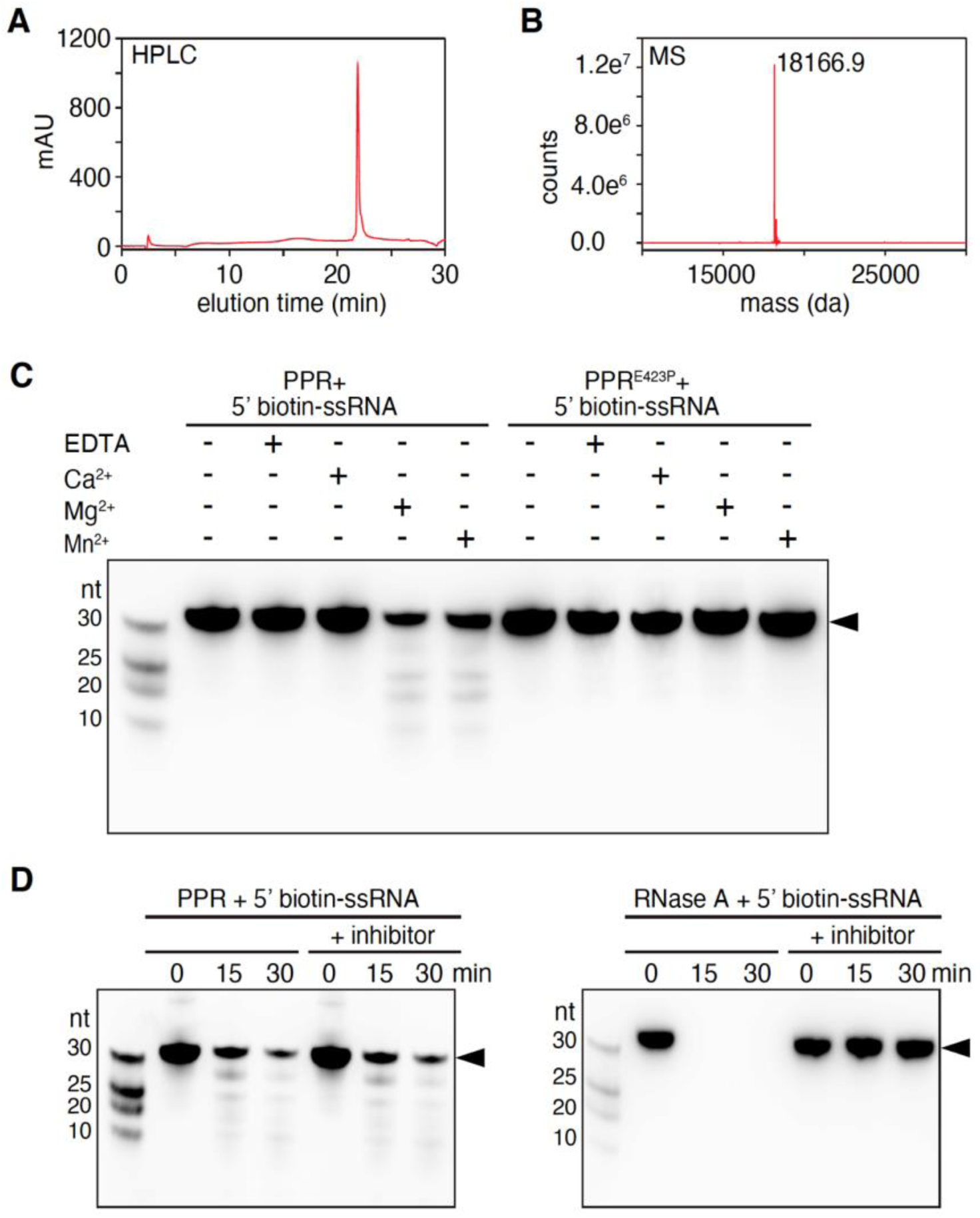
Purification and biochemical characterization of the PPR domain of POLRMT. (A) HPLC analysis of purified PPR domain using 210 nm UV absorbance. (B) Analysis of purified PPR domain by mass spectrometry, indicating a single peak at 18166.9 Dalton, which is the exact molecular weight of the recombinant PPR domain. (C) RNase activity of PPR domain requires divalent cations. Wild type PPR domain or PPR^E423P^ protein (0.5 µM) was incubated with the 5’-Biotin-labeled target ssRNA substrate (0.05 µM) in the presence of 10 mM EDTA, 5 mM Ca^2+^, 5 mM Mg^2+^, or 5 mM Mn^2+^ as indicated. Reaction was incubated for 15 min at 32 °C. Total reaction volume is 10 µl. RNA products were separated on a denaturing polyacrylamide gel (20%). Molecular size markers are indicated on the left. Arrowhead indicates the full-length RNA. (D) PPR protein is not sensitive to broad spectrum RNase inhibitor. Recombinant PPR domain (0.5 µM) was incubated with the 5’-Biotin-labeled ssRNA (0.05 µM) in the absence or the presence of RNase inhibitor after 15 and 30 min at 32 °C. RNase A was used as a control. RNA products were separated on a denaturing polyacrylamide gel (20%). Molecular size markers are indicated on the left. Arrowhead indicates the full-length RNA.

**Figure S3.**
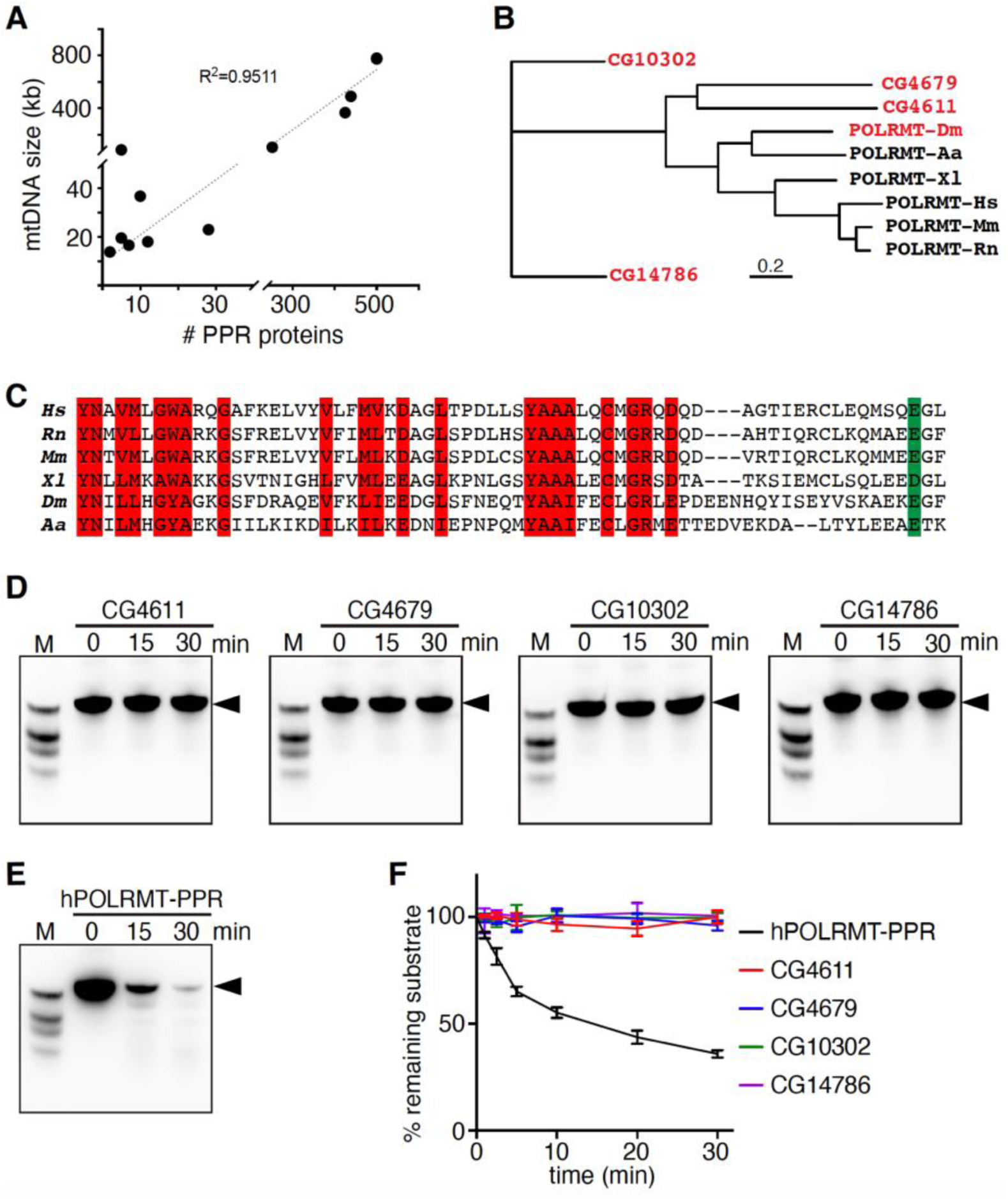
Evolution and phylogenic analyses of PPR proteins. (A) Correlation between the number of PPR genes and the mtDNA genome size in different organisms. R^2^, correlation coefficient. (B) Phylogenetic analysis of PPR domains in different PPR proteins using the neighbor-joining algorithm. The tree was constructed with PPR domains of *Drosophila melanogaster* CG10302, CG4679, CG4611, CG14786 and POLRMT of six different species. Distance scale, 0.2 (20%) divergence. (C) Sequence alignment of POLRMT PPR domains in *Drosophila melanogaster* (*Dm*) and five other species. Conserved residues are highlighted. (D) Four other PPR proteins from *Drosophila* do not have exoribonuclease activity. 5’-Biotin-labeled target ssRNA (0.05 µM) was incubated with CG4611, CG4679, CG10302 or CG14786 (0.5 µM) for indicated time (min) at 32 °C and the products were analyzed on a denaturing polyacrylamide gel (20%). Arrowhead indicates the full-length RNA. (E) PPR domain of human POLRMT has exoribonuclease activity. 5’-Biotin-labeled target ssRNA (0.05 µM) was incubated with recombinant PPR protein of human POLRMT (0.5 µM) for indicated time (min) at 32 °C and the products were analyzed on a denaturing polyacrylamide gel (20%). Arrowhead indicates the full-length RNA. (A) Pseudo-first-order cleavage kinetics of 5’-Biotin-labeled target ssRNA by PPR domain of human POLRMT (residues 218-368), CG4611, CG4679, CG10302 and CG14786. Each data point represents the average of three independent experiments. Error bars represent sd. Abbreviations in (B&C): Hs, *Homo sapiens;* Rn, *Rattus norvegicus*; Mm, *Mus musculus*; Xl, *Xenopus laevis*; Dm, *Drosophila melanogaster*; Aa, *Aedes aegypti*.

**Figure S4.**
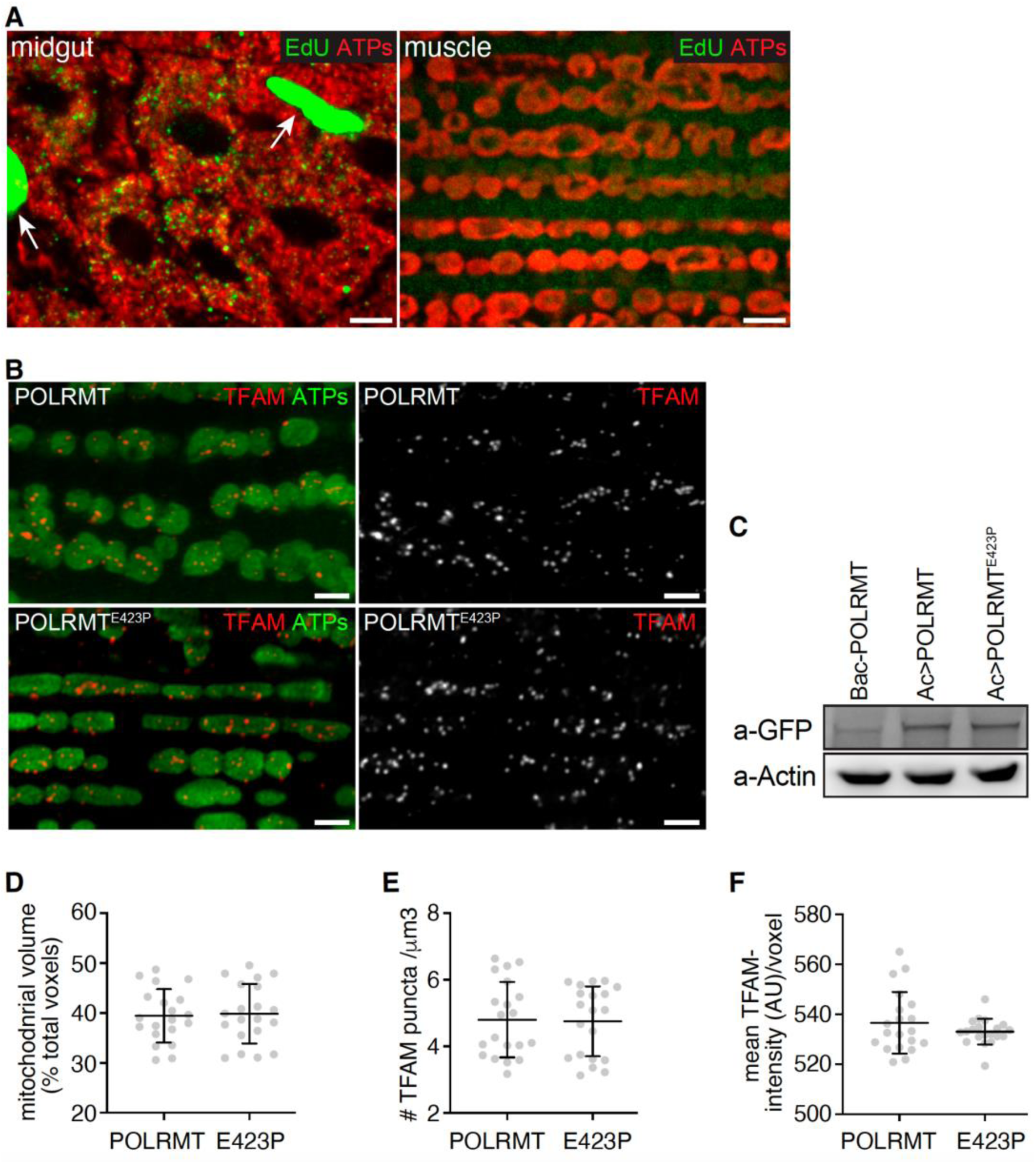
Overexpression of POLRMT or POLRMT^E423P^ does not affect mitochondrial nucleoids in adult flight muscles. (A) Representative images of *Drosophila* midgut (left panel) and indirect flight muscle (right panel) with EdU incorporation (Green) and ATP synthase (ATPs; Red) co-staining. Arrowheads indicate EdU incorporation into the nucleus of midgut cells. Scale bars, 10 μm. (B) Representative confocal images of indirect flight muscle from POLRMT and POLRMT^E423P^ expression flies stained for TFAM (Red) and ATP synthase (ATPs; Green). Scale bars, 10 μm. (C) Western blot showing that POLRMT and POLRMT^E423P^ overexpression levels (using ac-Gal4:PR driver) are increased compared to the POLRMT Bac clone expression. Actin is used as a loading control. (D) Quantification of mitochondrial contents by normalizing mitochondrial volume marked with ATP synthase (voxels) to total voxels of tissue in POLRMT or POLRMT^E423P^ expressing muscle (n=20). Error bars represent SD. p>0.05. (E) Quantification of TFAM puncta volume (voxels) per μm^3^ in indirect flight muscles (n=20). Error bars represent sd. p>0.05. (F) Quantification of TFAM mean intensity per voxel in indirect flight muscles (n=20). Error bars represent sd. p>0.05. Genotypes in (D-F) are: *w;;ac-Gal4:PR>UAS-POLRMT* (POLRMT) and *w*;;*ac-Gal4:PR*>*UAS-POLRMT^E423P^* (E423P).

**Figure S5.**
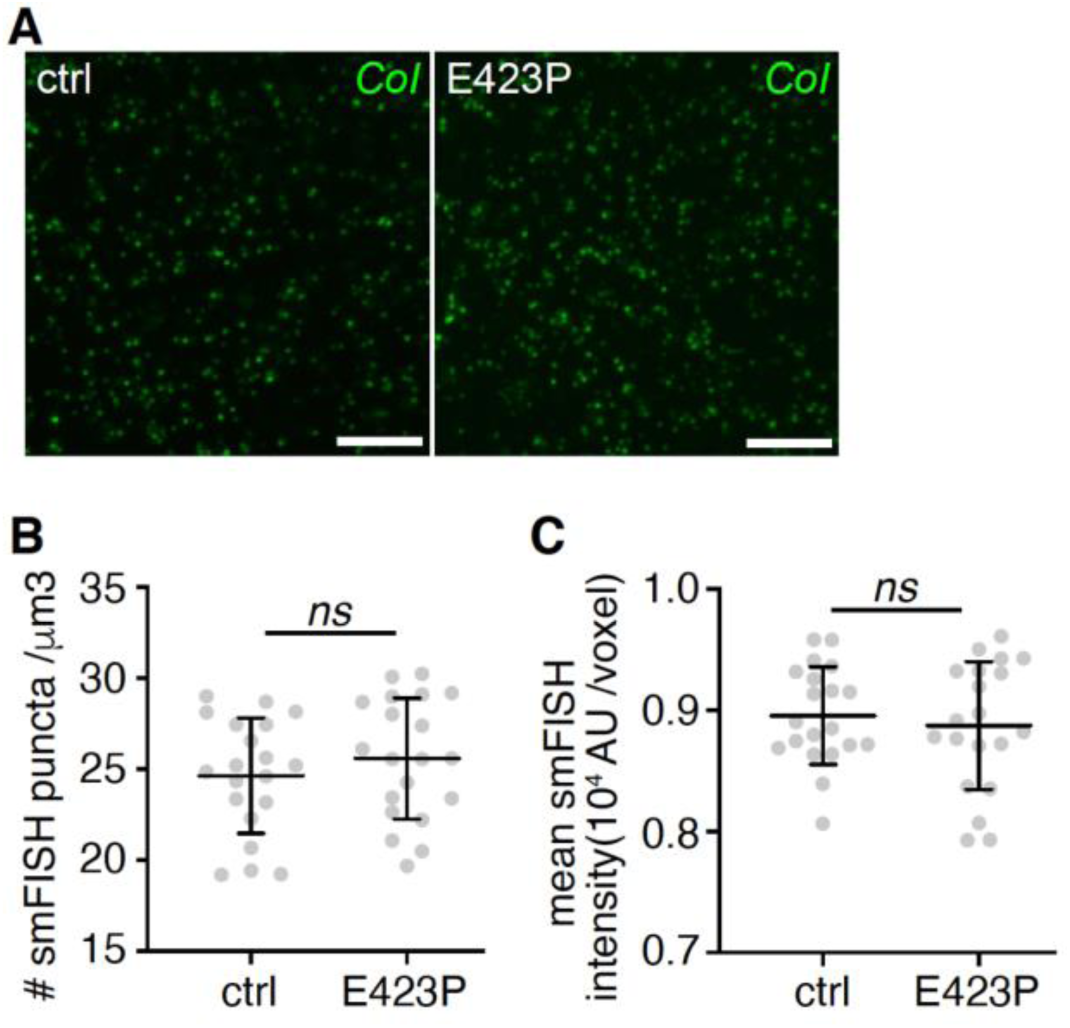
The ablation of POLRMT’s exoribonuclease activity does not affect mitochondrial transcription. (A) Visualization of the *mt: CoI* mRNAs in indirect flight muscles from ctrl and E423P flies by smFISH with RNA probes (Green). Scale bars, 10 μm. (B) Analysis of mitochondrial transcription by quantifying the number of smFISH puncta per μm^3^ in indirect flight muscle (n=20). Error bars represent sd. p>0.05. (C) Quantification of mitochondrial transcription illustrated as the mean smFISH intensity per voxel in indirect flight muscles (n=20). Error bars represent sd. p>0.05. Genotypes in (A-C) are: *w;;ac-Gal4:PR>UAS-POLRMT* (ctrl) and w;;ac-Gal4:PR>*UAS-POLRMT^E423P^* (E423P).

**Figure S6.**
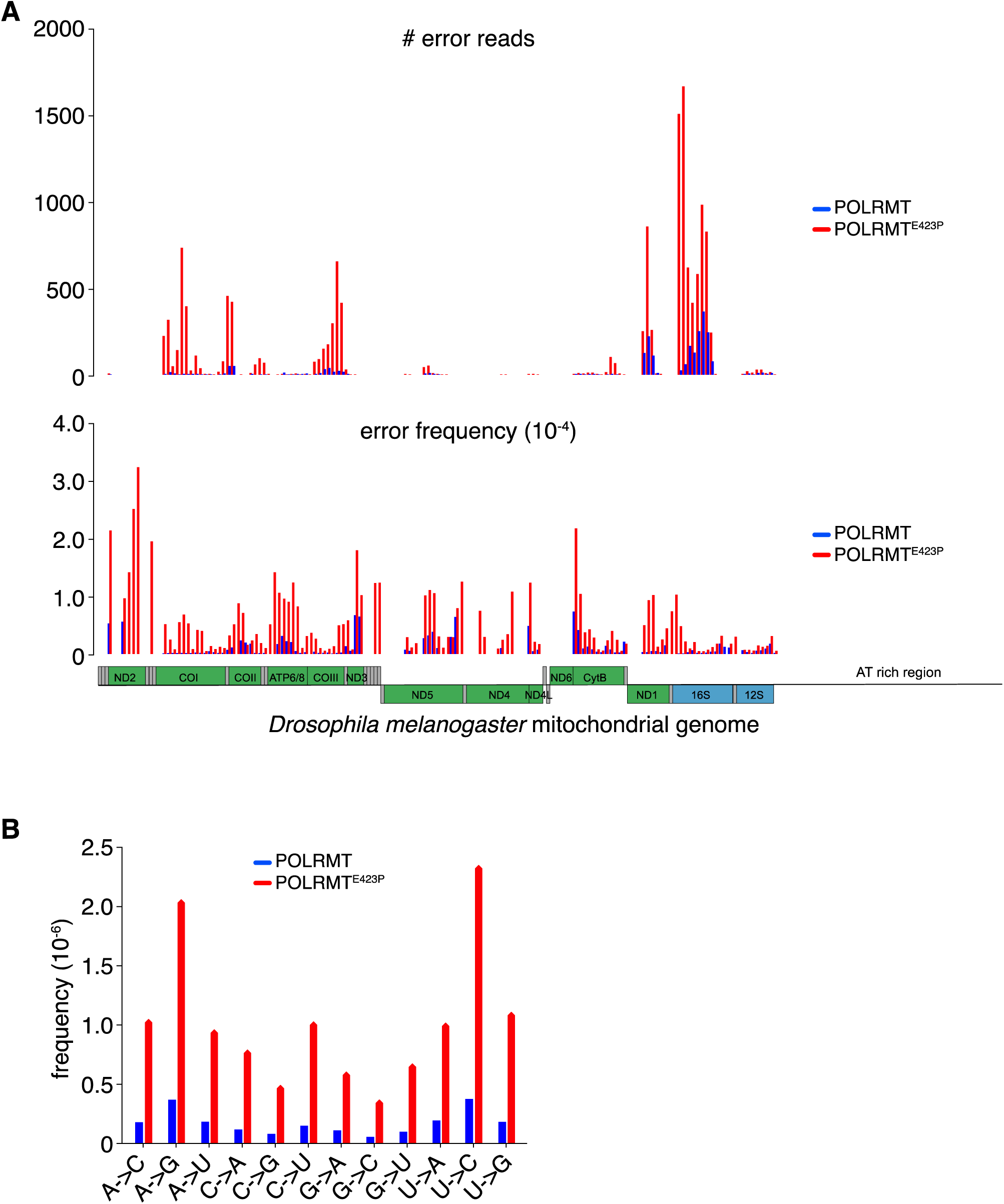
Overview of transcription errors in POLRMT and POLRMT^E423P^ overexpression flies. (A) The transcription errors detected from POLRMT and POLRMT^E423P^ overexpression flies were distributed randomly across the mitochondrial genome, and most errors were located in highly transcribed genes. “Error reads” indicates the total number of errors detected in a 100-bp bin. “Error frequency” was calculated by dividing the total number of errors reads by the total number of sequencing reads within that interval. (B) The spectrum of transcripts errors in POLRMT and POLRMT^E423P^ overexpression flies.

## Supplemental Table Legends

**Table S1. List of all unprocessed RNA frequency values of POLRMT and POLRMT^E423P^ overexpression flies, related to Figure 5**.

**Table S2. List of all tRNA counts in POLRMT and POLRMT^E423P^ overexpression flies, related to Figure 5**.

## Notes

https://data.mendeley.com/datasets/jsn8c49wpy/draft?a=1e8e1cc6-b2e9-4328-826b-1e94502d0c34

### Competing Interest Statement

The authors have declared no competing interest.

